# The Japanese wolf is most closely related to modern dogs and its ancestral genome has been widely inherited by dogs throughout East Eurasia

**DOI:** 10.1101/2021.10.10.463851

**Authors:** Jun Gojobori, Nami Arakawa, Xiaokaiti Xiayire, Yuki Matsumoto, Shuichi Matsumura, Hitomi Hongo, Naotaka Ishiguro, Yohey Terai

**Affiliations:** SOKENDAI (The Graduate University for Advanced Studies), Department of Evolutionary Studies of Biosystems, Shonan Village, Hayama, Kanagawa 240-0193, Japan; Research and Development Section, Anicom Specialty Medical Institute Inc., Chojamachi Yokohamashi-Nakaku, Kanagawaken 231-0033, Japan; Faculty of Applied Biological Sciences, Gifu University, Yanagido 1-1, Gifu 501-1193, Japan

## Abstract

The Japanese wolf (*Canis lupus hodophilax* Temminck, 1839) was a subspecies of the gray wolf that inhabited the Japanese Archipelago and became extinct 100-120 years ago. In this study, we determined the whole genomes of nine Japanese wolves from the 19^th^- early 20^th^ centuries and 11 Japanese dogs and analyzed them along with both modern and ancient wolves and dogs. Genomic analyses indicate that the Japanese wolf was a unique subspecies of the gray wolf that was genetically distinct from both modern and ancient gray wolves, lacking gene flow with other gray wolves. A Phylogenetic tree that minimizes the effects of introgression shows that Japanese wolves are closest to the dog monophyletic group among the gray wolves. Moreover, Japanese wolves show significant genetic affinities with East Eurasian dogs. We estimated the level of introgression from the ancestor of the Japanese wolves to the ancestor of East Eurasian dogs that had occurred in the transitional period from the Pleistocene to the Holocene, at an early stage after divergence from West Eurasian dog lineages. Because of this introgression, Japanese wolf ancestry has been inherited by many dogs through admixture between East Eurasian dog lineages. As a result of this heredity, up to 5.5% of modern dog genomes throughout East Eurasia are derived from Japanese wolf ancestry.

## Introduction

The phylogeny of gray wolves (*Canis lupus*) attracts wide attention from researchers and the public because wolves are the closest relatives to one of the most familiar animal species to humans, i.e., dogs. The extant gray wolves (*Canis lupus*) are divided into three lineages: the North American, Eurasian, and domestic dog lineages, including several now-extinct lineages that inhabited Eurasia during the Pleistocene (Ramos-Madrigal, et al. 2021). Recent phylogenomic analyses of gray wolves have shown that the North American gray wolf diverged at the basal ancestral position, followed by the Eurasian lineage (Fan, et al. 2016; Leathlobhair, et al. 2018). Dogs form a monophyletic clade which is the sister group to the Eurasian lineage of the gray wolf (Freedman, et al. 2014; Fan, et al. 2016; Leathlobhair, et al. 2018). Therefore, the hypothesis that the dog lineages have originated in Eurasia has been widely accepted. But there is still much debate concerning when, where, how many times, and from which population, the ancestor of dogs was domesticated (Leonard, et al. 2002; Savolainen, et al. 2002; Germonpré, et al. 2009; Pang, et al. 2009; Vonholdt, et al. 2010; Larson, et al. 2012; Axelsson, et al. 2013; Thalmann, et al. 2013; Freedman, et al. 2014; Perri 2016; Janssens, et al. 2018; Leathlobhair, et al. 2018; Perri, et al. 2021). Because no extant population of gray wolves has been reported to be more closely related to dogs than the other wolf populations, it is believed that the dog lineage has been domesticated from an extinct population of gray wolves (Larson, et al. 2012; Thalmann, et al. 2013; Freedman, et al. 2014; Skoglund, et al. 2015; Fan, et al. 2016; Frantz, et al. 2016). However, no information is available about this extinct population.

Many regions in Eurasia, including southern East Asia (Savolainen, et al. 2002; Pang, et al. 2009; Wang, et al. 2013; Wang, et al. 2016), Middle East (Vonholdt, et al. 2010), Central Asia (Shannon, et al. 2015), Europe (Thalmann, et al. 2013), and both West and East Eurasia (dual origin) (Frantz, et al. 2016), have been proposed as candidates for the origin of dogs, but the debate on the origin (single, or dual) as well as the timing of domestication still continues. Divergence between the Eurasian gray wolf and dog lineages has been estimated to be 20,000-40,000 years ago (Skoglund, et al. 2015; Botigué, et al. 2017). Based on phylogenomic analyses, dogs were initially reported to be genetically divided into two distinct lineages, i.e., the West and East Eurasian lineages (Freedman, et al. 2014; Shannon, et al. 2015; Frantz, et al. 2016; Botigué, et al. 2017; Leathlobhair, et al. 2018). Subsequent reports suggested an ancient divergence of the Arctic sled dog lineage (Sinding, et al. 2020), which is closely related to the pre-contact American dogs (Leathlobhair, et al. 2018). The phylogenetic relationship between the Arctic sled dog lineage and the West and East Eurasian lineages is conflicting (Larson, et al. 2012; Frantz, et al. 2016; Wang, et al. 2016), and these inconsistent topologies can be explained by either a high degree of admixture after the divergence of the three lineages, or by nearly simultaneous divergence (Frantz, et al. 2016; Zhang, et al. 2020). The West Eurasian and East Eurasian lineages diverged 17,000-24,000 years ago (Botigué, et al. 2017), and the Arctic sled dog lineage is estimated to have existed at least 9500 years ago (Sinding, et al. 2020).

Studies have suggested that wolf populations in Europe (Thalmann, et al. 2013), the Middle East (Vonholdt, et al. 2010), Central Asia (Shannon, et al. 2015), Siberia (Sinding, et al. 2020; Ramos-Madrigal, et al. 2021), and East Asia (Savolainen, et al. 2002; Wang, et al. 2016) have undergone introgression or bidirectional gene flow (Freedman, et al. 2014) with dogs. However, genomic introgression from gray wolves to dogs has been considered to be limited due to the presence of Eurasian wolves that do not show genetic affinity to any dog breed (Bergström, et al. 2020).

The Japanese wolf (*Canis lupus hodophilax* Temminck, 1839) was a subspecies of the gray wolf that inhabited Honshu, Shikoku, and Kyushu Islands in the Japanese Archipelago and became extinct 100-120 years ago (Ishiguro, et al. 2009). Molecular phylogenetic analysis of the mitochondrial genome suggests that the Japanese wolf diverged at the basal position of the extant gray wolf clade (Matsumura, et al. 2014; Matsumura, et al. 2020). Recent genome analysis of a “Honshu wolf” (one of the common names for the Japanese wolf) from the collection of the British Museum suggests that this individual is closely related to a lineage of Siberian wolves that existed in the Late Pleistocene and shows significant gene flow with Japanese dogs (Niemann, et al. 2021).

In this study, the genomes of nine Japanese wolves, including the type specimens, and 11 Japanese dogs were newly determined and analyzed. The analyses showed that 1) the Japanese wolf was a unique subspecies of the gray wolf that is genetically distinct from both extant and ancient gray wolves known to date, 2) the Japanese wolf is most closely related to a monophyletic group of dogs, and 3) Japanese wolf ancestry has introgressed into the ancestor of East Eurasian dogs at an early stage of their history after diverging from the West Eurasian lineages, and the genome derived from Japanese wolf ancestry has been inherited by many modern dogs, even in the West Eurasian lineages, through their historical admixture with East Eurasian lineages.

## Results

### Relationships between Japanese wolves and other dogs and gray wolves

In this study, original genomic DNA sequences of nine Japanese wolves (22-282 Gb) and 12 Japanese dogs (70-127 Gb) were determined (Table S1, Fig. S1). For the present study, we treated nine individuals with Japanese wolf type mitochondrial DNA haplotypes (Matsumura, et al. 2020) as the Japanese wolf. In addition, we used sequencing data of modern gray wolves with depth of coverage >20x in the database, ancient canids with relatively high coverage, and outgroup species from the database (Table S2). All sequence data were mapped to the reference genome sequence (CanFam3.1). After haplotype calling and gvcf file merging, <u>s</u>ingle <u>n</u>ucleotide <u>p</u>olymorphisms (SNPs) were genotyped. To examine the genetic relationship among the individuals used in this study, a principal component analysis (PCA) was performed using individuals with high coverage (Fig. 1A). In the PCA, Japanese wolves formed a distinct cluster, suggesting that Japanese wolves were genetically separated from dogs, gray wolves, and any of the outgroup species. The gray wolves were clustered into two groups, i.e., a North American/Arctic group and a Eurasian group along the PC1 axis. Dogs show an East-West cline along the PC2 axis. Dingoes and New Guinea singing dogs (NGSD) were the closest to Japanese wolves among dogs along the PC2 axis, followed by a cluster of Japanese dogs (Fig. 1A). Using the same data set we also generated an ADMIXTURE result with the lowest CV error at K=4 (Fig. 1B). In this analysis, Japanese wolves also formed clusters with higher K such as K=5 or K=6, indicating that their genetic composition was unique compared with that of the other dogs and gray wolves (Fig. 1B and S2).

**Figure 1.**
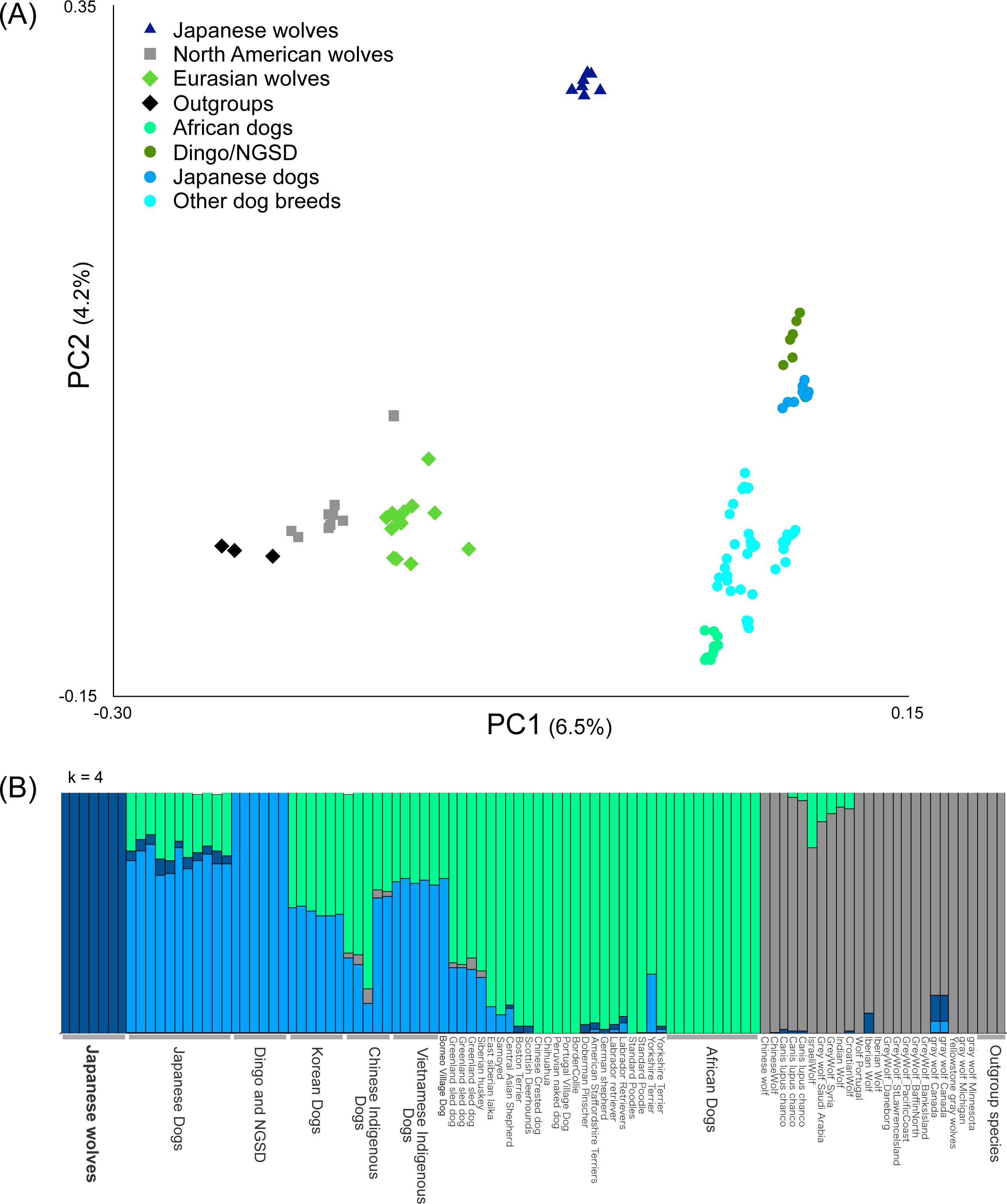
Relationships between Japanese wolves and other canids (A) Principal Components Analysis (PC1 versus PC2) of 100 samples based on 2,065,200 SNPs (see Table S2 for sample information). Colored circle, square, and triangle correspond to the names of dogs or wolves in the panel. (B) ADMIXTURE results based on SNP data for K = 4 (see Table S2 for sample information).

Next, we added the three Japanese wolf individuals with low coverage [Leiden b and c, (Jentink 1887), and a “Honshu wolf” (Niemann, et al. 2021), see Table S2] into the analysis. PCA showed that Leiden b and Honshu wolf were very close to the Japanese wolf cluster, while Leiden c was placed at an intermediate position between dogs and Japanese wolves (Fig. S3). ADMIXTURE analysis showed that Leiden b and Honshu wolf exhibit the same ancestry pattern as the other Japanese wolf individuals, while Leiden c seemed to be admixed with other dogs (Fig. S4). We used Patterson’s *f*4 statistic (Patterson, et al. 2012) to identify dog individuals with high genetic affinity to Leiden c to see which dog population was the source of the introgression. The dog that showed the highest affinity to Leiden c was the Japanese dog Shiba (Fig. S5A), and Leiden c contained 39% of the Shiba’s genome (Fig. S5B). In contrast, Leiden b showed no affinity with dogs (Fig. S5C). These results indicate that Leiden b and Honshu wolf are included in the group of Japanese wolves with a unique genetic composition, while Leiden c is a hybrid individual between Japanese wolves and dogs. Subsequently, we removed Leiden b, c and Honshu wolf individuals from further analyses due to their low coverage.

A previous study (Niemann, et al. 2021) suggested that Honshu wolf has a close relationship with a lineage of Siberian wolves that existed in the Late Pleistocene and was likely to be admixed with dogs. Therefore, we added the Pleistocene wolves to our dataset which are comprised of the modern gray wolves and dogs and performed PCA (Fig. S6). Pleistocene wolves were placed closely related to Eurasian wolves, while Japanese wolves formed a distinct cluster. Our ADMIXTURE analysis (Fig. S4) suggests that Honshu wolf does not contain more DNA components of dogs than the other Japanese wolf individuals. The differences between the results of Niemann et al. 2020 and this study could be caused by differences in the number of Japanese wolves used in the analyses and/or in the coverage of DNA sequencing.

### Phylogenetic position of the Japanese wolves

To determine the phylogenetic position of the Japanese wolves, a phylogenetic tree was constructed using the maximum likelihood (ML) method (Fig, 2A). Among gray wolves, North American/Arctic individuals branched off first at the basal position of the tree, followed by European/Middle Eastern and East Asian gray wolves (also see Fig. S7). Dogs formed a monophyletic clade including East Eurasian, West Eurasian, and sled dog lineages (Fig. 2A, S7), as shown in previous studies (Freedman, et al. 2014; Shannon, et al. 2015; Frantz, et al. 2016; Botigué, et al. 2017; Leathlobhair, et al. 2018; Sinding, et al. 2020). Japanese wolves formed a monophyletic clade that was a sister group to the monophyletic clade of dogs (Fig. 2A, S7). The sister group relationship between Japanese wolves and dogs was also supported by a tree inferred by SVDQuartets and a Neighbor-Joining tree based on identity-by-state (IBS) distance (Figs. S8, S9). Analysis using outgroup-*f3* statistics (Patterson, et al. 2012) also showed that the Japanese wolf was the most closely related to dogs among wolves (Fig. 2B). When we further divide the dogs into subpopulations, outgroup-*f3* statistics showed different results between dingo/NGSD and African dogs; dingo/NGSD was related most closely to Japanese wolf while African dog is related most closely to the Middle Eastern gray wolves (Fig. S10A and S10B). The different genetic affinities of dog populations to the Japanese wolf may have resulted from introgression between African dogs and Middle Eastern gray wolves (Vonholdt, et al. 2010; Bergström, et al. 2020).

**Figure 2.**
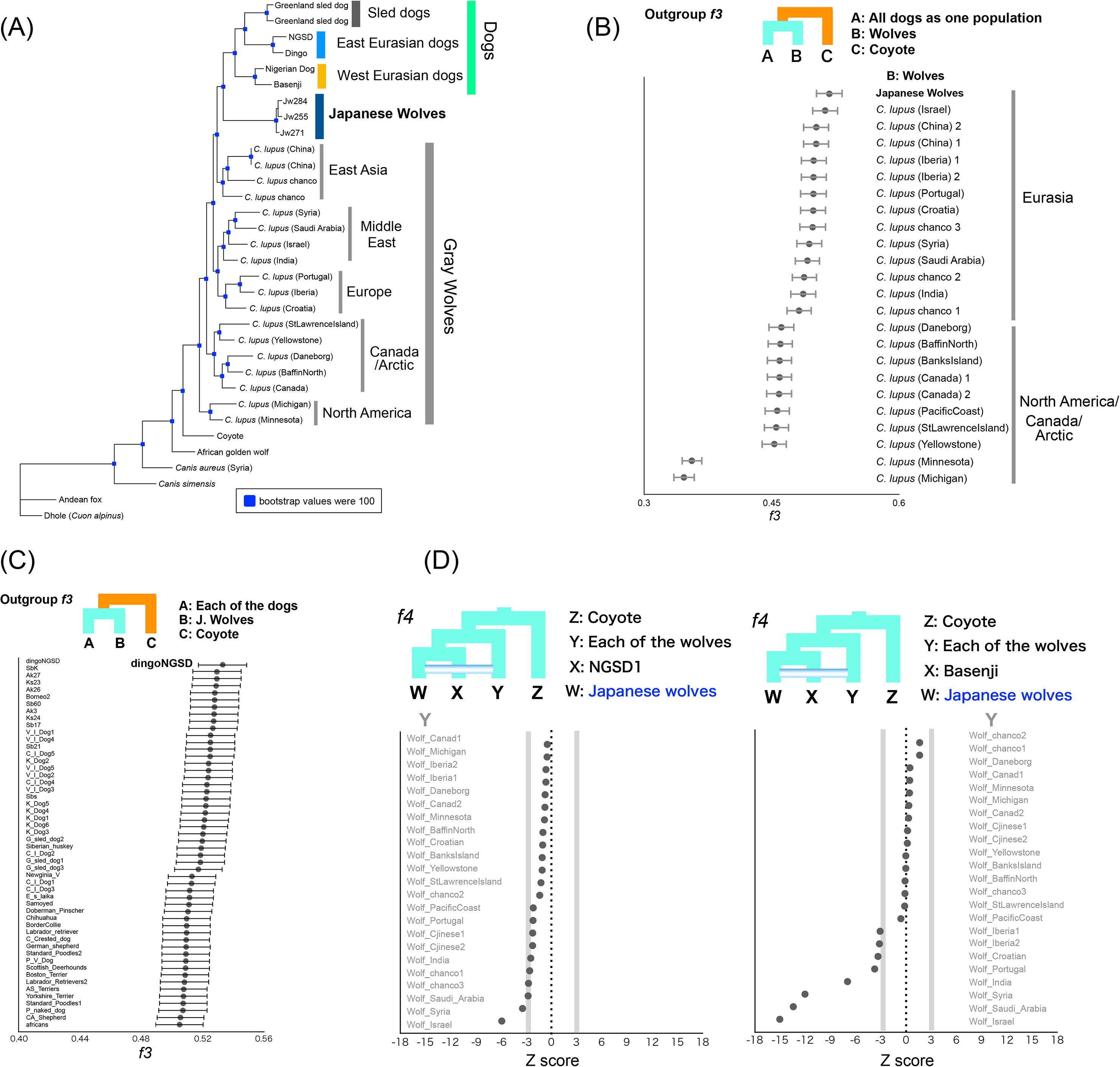
Phylogenetic relationships and genetic affinity between Japanese wolves and other canids. (A) Maximum likelihood tree based on 1,971,890 SNPs. Node labels shown in blue squares indicate bootstrap values out of 100. (B) Shared genetic drift between dogs and gray wolves measured by outgroup *f*3 statistics. Each of all dogs and Japanese wolves were used as populations. Each *f3* statistical value is plotted in order of highest to lowest value from the top, and the names of the wolves are shown on the right side of the panel. Error bars represent standard errors. (C) Shared genetic drift between Japanese wolf and all dogs measured by outgroup *f*3 statistics. Each of the African dogs and Dingo/NGSD individuals were used as populations. Each *f3* statistical value is plotted in order of highest to lowest value from the top, and the names of the dogs are shown on the left side of the panel. Error bars represent standard errors. (D) *f4* statistics testing the relationships between the Japanese wolf and all other wolves compared with NGSD1 (left panel) and Basenji (right panel). Each Z score is plotted in order of highest to lowest value from the top, and the names of wolves are shown on the left and right sides of the left and right panels, respectively. Gray lines show the Z score −3 and 3.

Since the tree topology in phylogenetic analyses could be affected by introgression between taxa, a phylogenetic tree using taxa showing minimal introgression effects is expected to be the most accurate representation of population branching. Therefore, in order to obtain such a tree, we examined introgression between Japanese wolves and other dogs and wolves. We compared the genetic affinity of each dog with the Japanese wolves using *f3* and *f4* statistics, and found that dogs of the East Eurasian lineage (Fig. S7), in particular, dingoes, NGSDs, and Japanese dogs, showed significant affinity with the Japanese wolves (Z score > 3) (Fig. 2C, S11). In contrast, dogs of the west Eurasian lineage, in particular dogs from Africa, showed low affinity to Japanese wolves (Fig. 2C, S11). *f4* statistics showed no affinity between any of the gray wolf populations and Japanese wolves (Fig. 2D). Possibilities of gene flow between gray wolves except for the Japanese wolf and dogs were also examined using *f4* statistics. Gray wolves in the Middle East showed strong affinity with dogs (Fig. S12), consistent with previous reports (Vonholdt, et al. 2010; Bergström, et al. 2020). Based on these results, we reperformed phylogenetic analysis to confirm the relationship between Japanese wolves and dogs. To minimize the effect of introgression between wolves and dogs, we included African dogs as the sole representatives of dogs, and excluded gray wolves from the Middle East. Even in the phylogenetic tree obtained from this analysis, the Japanese wolf still formed a sister clade with African dogs (Fig. S13). Thus, we concluded that the most closely related wolves to the dog lineage are the Japanese wolves.

### The genome of the Japanese wolf ancestor in the dog genome

Japanese wolves showed strong affinity with many East Eurasian dogs (*f3*, *f4* statistics) (Figs. 2C, S11), which may be caused by the introgression of dog genomes into Japanese wolves (Japanese wolf or its ancestors, hereafter simply referred to as the “the Japanese wolf genome”) or vice versa. A recent report suggested widespread gene flow from dogs to gray wolves, but little gene flow from gray wolves to dogs, based on the existence of gray wolves that have no affinity with dogs (Bergström, et al. 2020). We investigated the direction of gene flow between Japanese wolves and East Eurasian dogs using the *f4*-ratio (Patterson, et al. 2012). We found that the degree of genome introgression from the Japanese wolf lineage to dogs was the highest in dingoes and NGSDs (5.5%) followed by Japanese dogs (3-4%), as well as in dogs of other East Eurasian lineages (Fig. 3A). In contrast, genomic introgression from dogs to the Japanese wolf genome was not supported (Fig. S14). We further analyzed the possibility of a small proportion of genomic introgression from dogs to the Japanese wolf genome by the *f4* statistics. If the dog genome is introgressed into the Japanese wolf genome, the genetic affinities between the Japanese wolf and dogs would be different between individuals. Indeed, the degree of affinity of Japanese wolves with dongo/NGSD varied among individuals (Fig. S15) and Jw258, Jw269, and Jw271 showed significant affinity to Japanese dogs (Fig. S16). This difference in affinity suggests that the Japanese wolf genome contains a small proportion of the dog genome that is undetectable by *f*4-ratio.

**Figure 3.**
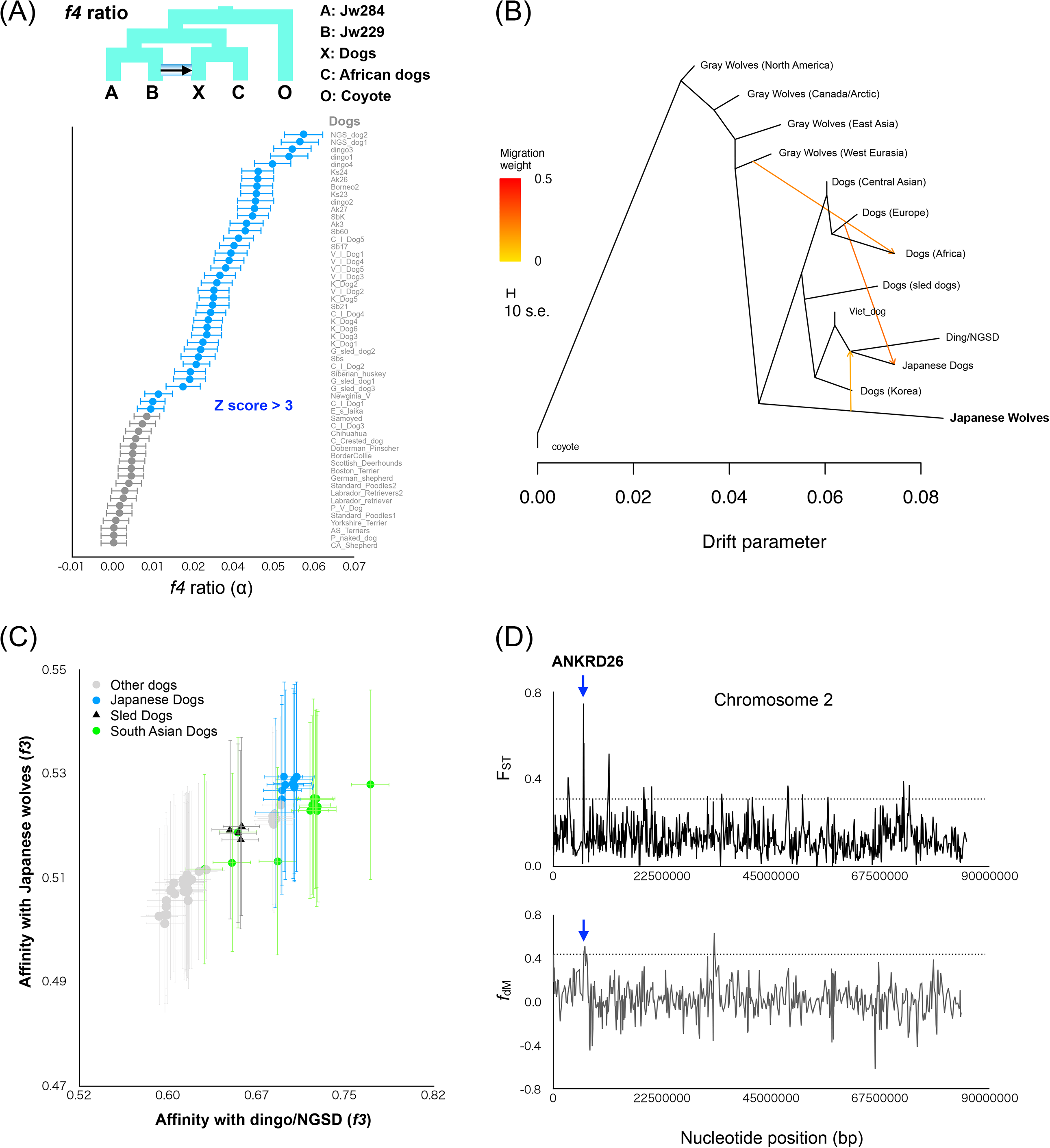
Admixture between Japanese wolves and the other canids. (A) *f4*-ratio test to estimate proportion of genome introgression from the Japanese wolf to dogs. Each *f4*-ratio α value is plotted in order of highest to lowest value from the top, and the names of the dogs are shown on the right side of the panel. Error bars represent standard errors. Z score above 3 is colored in blue. (B) TreeMix admixture graph built using LD-pruned data (150,502 sites) on a dataset consisting of 88 dogs/wolves merged into 13 groups according to their phylogenetic relationships. (C) *f3* statistics testing whether dogs share more alleles with dingo/ NGSD (x-axis) or Japanese wolf (y-axis). Dots show the *f3* statistics, and horizontal and vertical error bars represent standard errors for tests with the African dogs (x-axis) and dingo/NGSD (y-axis), respectively. Each of the Japanese wolves and dingo/NGSD individuals were used as populations. (D) Sliding window analyses of the F_ST_ values (y-axis: upper panel) and *f*_dM_ (y-axis: lower panel) in windows of 50 SNPs using a 25 SNPs slide across chromosome (x-axis). Dashed lines show the 99th percentiles. Blue arrow indicates overlapping regions above 99th percentiles between upper and lower panels. The gene name above the arrow is the gene in the overlapped region.

The degree of genomic introgression from Japanese wolves to dogs was higher in East Eurasian than in West Eurasian dogs. It also varied among the dogs of East Eurasia. This variation may have been caused by multiple introgression events between the ancestors of Japanese wolves and dogs in different regions, or by a single introgression followed by diffusion of the Japanese wolf genome into various dog populations. To determine which hypothesis is more likely, we first examined the degree of gene flow among dogs in different regions. African dogs and dingo/NGSD represent opposite edges of the dog cluster in the PCA (Fig. 1A), and show the lowest and highest affinities with Japanese wolves, respectively (Fig. 2C). Among dogs, African dogs show the lowest affinity with dingo/NGSD, while dingo/NGSDs show the lowest affinity with African dogs (Fig. S17, S18). Dingoes are estimated to have been introduced to Australia between 4600 and 18300 years ago based on their mitochondrial DNA (Oskarsson, et al. 2012), with archaeological evidence supporting the arrival of at least 3500 years ago (Milham and Thompson 1976). It is considered that the dingoes have been isolated in Australia since then (Larson, et al. 2012; Fan, et al. 2016). African dogs are estimated to have migrated to Africa 14,000 years ago (Liu, et al. 2018) and have been isolated since then (Larson, et al. 2012; Fan, et al. 2016). The African Dog and Dingo/NGSD are included in the West Eurasian and East Eurasian lineages, respectively (Fig. S7). Therefore, they are likely to be the oldest dogs of their respective lineages. The *f4* statistics biplot showed that dogs showing higher affinity with dingo/NGSD show lower affinity with African dogs while dogs showing higher affinity with African dogs show lower affinity with Dingo/NGSD (Fig. S19). This negative correlation suggests most of dog populations were formed through extensive past mixing between East and West Eurasian lineages represented by dingo/NGSD and African dogs, respectively. Indeed, several dogs in South and East Asia are genomically characterized as dingo/NGSD admixed with African dogs by negative values of *f3* statistics (Patterson, et al. 2012) (Fig. S20).

Next, we examined the degree of introgression between dogs from different regions and Japanese wolves. TreeMix analysis indicates an introgression from the ancestor of Japanese wolves into the common ancestor of dingo/NGSD and Japanese dogs (Fig. 3B). The *f3* biplot of affinities with dingo/NGSD and with Japanese wolves shows a positive correlation among dogs, indicating that the Japanese wolf genome has become widespread through the admixture between West and East Eurasian dog lineages and persists in the modern genomes of the East Eurasian lineage (Fig. 3C). Therefore, it is likely that the genome of the Japanese wolf ancestor was introgressed into an ancestral lineage of the East Eurasian lineage after the split of West and East Eurasian lineages. Subsequently, the East Eurasian lineage containing the Japanese wolf ancestry admixed with the West Eurasian lineage, resulting in differences in affinities with the Japanese wolves. The Southeast Asian dogs (Fig. 3C green) have a relatively higher affinity with Dingo/NGSD compared to their affinity with Japanese wolves, suggesting gene flow between the Southeast Asian dogs and dogs that carried no Japanese wolf genome after the first admixture event. Similarly, Japanese dogs (Fig. 3C blue) have a relatively higher affinity with Japanese wolves compared to their affinity with Dingo/NGSD, suggesting subsequent gene flow between Japanese dogs and Japanese wolves. Hence, it is likely that the difference in the degree of genomic introgression from Japanese wolves to dogs was caused by a single introgression followed by diffusion of the Japanese wolf genome into various dog populations.

### Genomic regions derived from Japanese wolf that contribute to the traits of Japanese dogs

The results of *f4*-ratio analysis showed that Akita, Shiba, and Kishu dogs contain 36-45% West Eurasian dog genomes (Table S3). It is expected that the genomic regions containing genes responsible for traits of the Japanese dogs are highly differentiated from those of West Eurasian dogs. If such regions overlap with the Japanese wolf-derived genomic regions (3-4%) in the modern Japanese dog genome, it could be that Japanese wolf-derived genes are responsible for the characteristic traits of the Japanese dog. To investigate such regions, we examined genomic regions derived from Japanese wolves using a *f*_dM_ sliding window and regions differentiated between Japanese dogs and West Eurasian dogs using an F_ST_ sliding window, and extracted overlapping regions. Six regions were found, which included four genes and the upstream or downstream regions of four genes (Fig. 3D, Fig. S21, Table S5). These regions may be candidates of genomic regions derived from Japanese wolf that contribute to the traits of Japanese dogs.

## Discussion

The Japanese wolves are likely to have been isolated in the Japanese archipelago until their extinction only 100 years ago. This study reveals that they form a monophyletic group with no evidence for gene flow with other Eurasian gray wolves.

One notable aspect of Japanese wolves is their phylogenetic position. In Eurasia, our phylogenetic analysis showed that the European lineage of the gray wolves diverged at the basal position, followed by the Middle East and East Asian lineages. In the East Asian lineage, the monophyletic group of Japanese wolves and the dog lineage form a sister-group relationship. The order in which Eurasian wolf lineages diverged is from west to east in geographical order on the Eurasian continent. Considering these phylogenetic and geographic relationships, it is most likely that it was in East Asia that the divergence between the Japanese wolf and the dog lineages has occurred. In other words, the extinct population of the gray wolf from which dogs are suspected to have been domesticated (Larson, et al. 2012; Thalmann, et al. 2013; Freedman, et al. 2014; Skoglund, et al. 2015; Fan, et al. 2016; Frantz, et al. 2016) is closely related to the ancestor of the Japanese wolf and is likely to inhabit East Asia. This hypothesis does not directly imply that the origin of dog domestication was in East Asia, nor does it directly imply that the earliest dogs descended from the ancestor of Japanese wolves. Although the domestication process would have been initiated with the animals’ association with humans (Larson, et al. 2012; Perri 2016; Janssens, et al. 2018), our phylogenetic analyses provide no evidence for when dog lineages began to associate with humans. Further archaeological evidence in the studies of ancient “proto-dog” populations are required to clarify the beginnings of the dog-human relationship.

This study suggests ancient genomic introgression from the Japanese wolf to dogs, most likely to the ancestor of the East Eurasian lineage. The divergence between the dog lineage and the Eurasian gray wolves has been estimated to be 20,000-40,000 years ago (Skoglund, et al. 2015; Botigué, et al. 2017). Dogs have been reported to have split into West Eurasian, East Eurasian, and sled dog lineages in their early divergence (Freedman, et al. 2014; Shannon, et al. 2015; Frantz, et al. 2016; Botigué, et al. 2017; Leathlobhair, et al. 2018). Because a 9500-year-old sled dog (Sinding, et al. 2020) already contained the same proportion of the Japanese wolf genome as the modern sled dog (2%: Table S4), the genomic introgression of the ancestor of the Japanese wolf to the East Eurasian lineage of dogs must have occurred before the establishment of the sled dog lineage at least 9500 years ago during the transitional period from the Pleistocene to the Holocene and shortly after the divergence of the East and West Eurasian dog lineages. The genome of NGSD was estimated to contain the Japanese wolf genome (5.5%). It is estimated that the NGSD lineage already existed by 10,900 years ago (Bergström, et al. 2020), which also supports the hypothesis that the introgression from the ancestor of Japanese wolves into dogs had occurred in the Pleistocene. The ancient dog genome data from two European individuals (4,800 and 7,000 years ago) already contained about 1.6% of the Japanese wolf genome (Table S4). Since the gene flow from the Southeast Asian dog ancestry to the ancestor of these two ancient European dogs has been reported (Botigué, et al. 2017), the genome of the Japanese wolf may have been transmitted to European dogs via the Southeast Asian dog ancestry more than 7,000 years ago. Although the Japanese wolf has only ever been found living in the Japanese archipelago, it is unlikely that the introgression between the ancestor of the Japanese wolf and dogs of the East Eurasian lineage occurred in the Japanese archipelago. An ancient sled dog (Sinding, et al. 2020) excavated at the same time as the excavation of the oldest dog in Japan (9600 years ago) (Shigehara and Hongo 2000) already possessed the Japanese wolf genome, suggesting that introgression between the ancestor of Japanese wolves and dogs of the East Eurasian lineage had occurred before dogs were brought to the Japanese archipelago. Therefore, the introgression between the ancestral Japanese wolf and the East Eurasian lineage of dogs is most likely to have occurred somewhere in East Asia.

The dogs with a high proportion of the Japanese wolf genome are the dingo/NGSD (5.5%) and the Japanese dogs (3-4%). The high proportion of the dingo/NGSD is inferred to be due to their isolation in the islands of Southeast Asia and Australia, where they have escaped admixture with the West Eurasian dog lineage. NGSD is estimated to be an admixture of two lineages (Bergström, et al. 2020), and thus one of the admixed lineages may have possessed a higher proportion (∼10%) of the Japanese wolf genome than NGSD. In contrast, Japanese dogs contain about 36-45% of the West Eurasian genome (Table S3), even though a high proportion of the Japanese wolf genome persists in their genomes. After the first introgression with the East Eurasian dog ancestry, the Japanese wolf genome may have introgressed into the Japanese dog genomes in the Japanese archipelago. This hypothesis is supported by the ratio of affinity with Japanese wolves and with dingoes/NGSDs, which tends to be higher in Japanese dogs (Fig. 3C). In addition, the Japanese dogs have the highest affinity to the Japanese wolf among all dogs (Fig. S16), suggesting that the Japanese wolf genome was recently introgressed into the Japanese dog genome.

Although only a small proportion of the Japanese wolf genome persists in modern dog genomes, the Japanese wolf genome might have an effect on dog characteristic traits. We searched for genomic regions in Japanese dogs that derived from the Japanese wolf, and that were highly differentiated between Japanese dogs and West Eurasian dogs. Six regions were found with these criteria (Fig. 3D, Fig. S21, Table S5). Despite the gene flow from Eurasian dogs, these regions were differentiated between the genomes of Japanese and West Eurasian dogs. Therefore, these regions are expected to have been selected in the Japanese dogs and are the candidates of genomic regions potentially responsible for phenotypic characteristics of Japanese dogs. Further analysis of the regions selected in East Eurasian dogs and a genome-wide association study for East Eurasian dog traits should reveal the effects of the Japanese wolf genome on dog traits.

In this study, we demonstrated that the Japanese wolf is a sister group with the monophyletic clade of dogs. Our original results support the hypothesis that the modern dog lineage was domesticated from an extinct population of gray wolves (Larson, et al. 2012; Thalmann, et al. 2013; Freedman, et al. 2014; Skoglund, et al. 2015; Fan, et al. 2016; Frantz, et al. 2016), and the Japanese wolf is the closest to this now-extinct gray wolf population. In addition, we estimated the levels of introgression from the ancestor of Japanese wolves to the ancestor of East Eurasian dogs. Accordingly, the Japanese wolf genome is expected to be involved in the early stages of dog domestication. Further analysis of the genome of the Japanese wolf and ancient dog genomes, in particular from East Eurasia, will continue to shed light on the origins of dog domestication.

## Materials and methods

### Samples, DNA extraction, and sequencing

Japanese Wolf and Japanese dog DNAs were extracted and used in our previous studies (Matsumura, et al. 2014; Matsumura, et al. 2020). The sample locations are listed in Table S1. DNAs of two individuals of Shiba (Shiba_shiro and Shiba_kuro) were extracted from blood samples using a DNeasy Blood & Tissue Kit (Qiagen) following the manufacturer’s instructions. The NEBNext Ultra II DNA Library Prep Kit for Illumina (New England Bio Labs, Ipswich, MA, USA) was used to construct libraries from genomic DNA following the manufacturer’s instructions. Paired-end (2 × 150 bp) sequencing was performed on the Illumina HiSeq X or NovaSeq 6000 platforms.

For Leiden b and Leiden c, the genome capture was performed using the SeqCap EZ Hybridization and Wash Kit (Roche, Basel, Switzerland), SeqCap EZ Accessory Kit v2 (Roche), SeqCap HE-Oligo Kit (Roche), and SeqCap EZ Pure Capture Bead Kit (Roche) following the manufacturer’s instructions for SeqCap EZ Library SR (Roche), with minor modifications. Briefly, Biotin-labeled genomic DNA fragments from Shiba were used as hybridization probes, instead of the SeqCap EZ library (Roche). Leiden b and Leiden c libraries were mixed with 135 ng of Biotin-labeled genomic DNA fragments and were hybridized at 47°C for 72 h. Other procedures were performed in accordance with the manufacturer’s instructions. Total mapped reads are listed in Table S1.

### Extraction of SNPs and vcf file preparation

Sequence reads from the genomic DNA libraries of nine Japanese wolves, eleven Japanese dogs (Table S1) as well as 88 wolves/dogs and six outgroup species from the database (Table S2) were trimmed to remove adaptor sequences and mapped to the dog reference genome (CanFam3.1) using CLC Genomics Workbench (https://www.qiagenbioinformatics.com/). Reads showing high similarity (> 90% in > 90% of read length) were mapped to the reference genome sequences and reads mapped to more than one position were removed (“ignore” option for reads mapped to multiple positions). The mapping data was exported in bam file format and sorted and indexed using samtools (Li, et al. 2009). The duplicated reads in bam files were marked by the MarkDuplicates algorithm implemented in GATK v4.2 (https://gatk.broadinstitute.org/hc/en-us). We performed genotype calling on all individuals analyzed in this study using the HaplotypeCaller algorithm in GATK v4.2. Genotypes of all individuals were output as gvcf format (-ERC GVCF option) and filtered by the VariantFiltration algorithm in GATK v4.2 with default parameters. All gvcf files were combined into a single gvcf format file by the CombineGVCFs algorithm in GATK v4.2. The combined file was genotyped by the GenotypeGVCFs algorithm and filtered by Filtervcf in GATK v4.2 with parameters; --filter-expression “QD < 2.0” --filter-name “QD2” --filter-expression “QUAL < 30.0” --filter-name “QUAL30” --filter-expression “FS > 200.0” --filter-name “FS200” --filter-expression “SOR > 10.0” -filter-name “SOR10” --filter-expression “ReadPosRankSum < −20.0” --filter-name “ReadPosRankSum-20”.

To maximize the number of SNPs for analyses without missingness, we prepared vcf files for each analysis using vcftools (Danecek, et al. 2011). When including all individuals except for ancient samples and Leiden b, Leiden c, and Honshu wolf (see the sample combination in Table S2), we removed all indels, singleton, and doubleton sites to eliminate PCR and sequencing errors that may have occurred in one individual, and extracted bi-allelic sites with coverage equal to or more than three in all individuals and with GQ values equal to or more than eight in all individuals. Mutations due to DNA damage at both ends of fragments were less than 1% in Japanese wolves (Fig. S1), therefore we can infer that mutations by DNA damage in the sequences of Japanese wolves were removed by this filtration. For analyses with ancient samples, sites were filtered using the same conditions as with modern samples. In addition, we used only transversion sites for the analyses of ancient samples. For the PCA and ADMIXTURE analyses with Liden b and Liden c, sites with missingness greater than 3% and minor allele frequency (MAF) < 0.04 were excluded. Sites were filtered using the same conditions as with modern samples, and only transversion sites were used.

### Phylogenetic analysis

The SNPs in a vcf file including dogs, wolves and Japanese wolves (see Table S2) were converted to PHYLIP format. 10 kb sequences from the 5’ end of the PHYLIP format file were extracted and a model for Maximum Likelihood method was selected using MEGA ver. X (Kumar, et al. 2018). A phylogenetic tree was constructed using the Maximum Likelihood (ML) method using PhyML ver. 3.2 (Guindon, et al. 2010) with a model selection option “-m GTR” and with 100 bootstrap replications. ML trees were constructed using all individuals (Fig. S7: 489,524 SNPs), selected individuals (Fig. 2A: 1,971,890 SNPs), and outgroup species (Table S2), wolves, Japanese wolves, and African dogs (Fig. S13: 2,065,200 SNPs).

The same vcf file used in the ML method (Fig. S7: 489,524 SNPs) was converted to NEXUS format. A phylogenetic tree was constructed by the svdq algorithm (Chifman and Kubatko 2014) in PAUP* ver. 4a (Wilgenbusch and Swofford 2003) with 100 bootstrap replications (Fig. S8). We used PLINK ver. 1.9 (Purcell, et al. 2007) with an option “—distance 1-ibs” to calculate an Identity By State (IBS) distance matrix using 1,992,260 SNPs. A neighbour joining tree was constructed from the IBS distance matrix using MEGA ver. X (Kumar, et al. 2018)(Fig. S9).

### Principal component analysis and ADMIXTURE

We performed a principal component analysis (PCA) using PLINK ver. 1.9 (Purcell, et al. 2007) with an option “--indep-pairwise 50 10 0.1” to explore the affinity among gray wolves, Japanese wolves, and dogs (Figure 1A). We also performed PCA with type specimens of Japanese wolf (Fig. S3), and with ancient canids (Fig. S6).

ADMIXTURE ver. 1.3 (Alexander and Lange 2011) was run on the dataset of modern samples (Fig. 1B and Fig. S2) and modern specimens with type specimens of Japanese wolf (Fig. S4) assuming 2 to 8 clusters (K=2-8).

### *f3*, *f4* statistics, and *f4*-ratio

*f*3, *f4* statistics, and *f4*-ratio implemented in ADMIXTOOLS ver. 7.0.1 (Patterson, et al. 2012) were used to evaluate the shared genetic drift among gray wolves, Japanese wolves, and dogs. We used the same SNPs data set used for the phylogenetic analysis for all modern samples (489,524 SNPs). For Liden b and Leiden c analyses, vcf files were prepared to maximize the number of SNPs (Table S2).

Outgroup *f3* statistics were calculated to explore shared genetic drift between all dogs and each of the wolves (Fig. 2B), African dogs and each of the wolves (Fig. S10A), dingo/NGSD dogs and each of the wolves (Fig. S10B), Japanese wolf and each of the dogs (Fig. 2C), African dogs and each of the other dogs (Fig. S17A), and dingo/NGSD dogs and each of the other dogs (Fig. S17B). *f3* statistics were calculated to test the genomic mixture of African and dingo/NGSD dogs in all dogs (Fig. S20).

*f*4 statistics were calculated to explore shared genetic drift between each of the dogs and Leiden c (Fig. S5A: 38,254 SNPs), each of the dogs and Leiden b (Fig. S5C: 83,259 SNPs), each of the wolves and Japanese wolves (Fig. 2D), each of the dogs and Japanese wolves (Fig. S11), each of the dogs and each of the wolves (Fig. S12), dingo/NGSD and Japanese wolves (Fig. S15), each of the dogs and Japanese wolves (Fig. S16), NGSD1 or Basenji and each of the dogs (Fig. S18), and dingoes and each of the other dogs, and African dogs and each of the other dogs (Fig. S19)

### TreeMix

To examine the admixture events, we used TreeMix (Pickrell and Pritchard 2012) to build a tree with admixture edges. We used major groups of gray wolves and dogs as follows; Gray Wolves (North America: n = 2), Gray Wolves (Canada/Arctic: n = 8), Gray Wolves (East Asia: n =5), Gray Wolves (West Eurasia: n = 7), Japanese Wolves (Japan: n = 7), Dogs (Central Asian: n = 4), Dogs (Europe: n = 11), Dogs (Africa: n = 10), Dogs (sled dogs: n = 4), Dogs (Vietnamese Indigenous: n = 5), Ding/NGSD (Oceania: n = 7), Japanese Dogs (Japan: n = 11), and Dogs (Korea: n = 6). The SNPs were pruned based on linkage disequilibrium (LD-pruning) by using plink with an option “--indep-pairwise 50 10 0.12. As a result of LD-pruning, 150,502 SNPs were used for TreeMix.

### Assessing DNA damage patterns

We used mapDamage ver. 2.2.0 (Ginolhac, et al. 2011) to assess DNA damage patterns in the Japanese wolf samples sequenced in this study. Mapped reads from the Japanese wolf samples showed slightly increased proportion (equal to or less than 1%) of C to T and G to A substitutions at the 5’ and 3’ read ends, respectively (Fig. S1).

### Calculation of maximum contamination rate

We used substitutions in mitochondria DNA specific to Japanese wolf to assess the contamination rate of the other animal DNA in Japanese wolf DNA. Fifteen fixed substitutions unique to Japanese wolf were selected using an alignment of mitochondria DNA sequences including gray wolves, Japanese wolves, and dogs used in previous studies (Matsumura, et al. 2014; Matsumura, et al. 2020). The lowest coverage at fifteen sites was 48 (highest was 28,324) in the mapping result to the mitochondria genome in CanFam3.1. We calculated the average mapping ratio in fifteen sites. The ratio of the reads mapped to the fifteen sites without substitutions specific to Japanese wolf were assumed as the maximum contamination rate (Table S6), because the mitochondria DNA like sequences are found in the nuclear genome.

### Detection of genomic regions that are derived from Japanese wolf and may have contributed to Japanese dog characteristics

The genomic regions containing genes that form the characteristics of the Japanese dog are expected to be largely differentiated between the Japanese dog and the West Eurasian dog. To detect such regions, we examined sequences differentiated between Japanese dogs and West Eurasian dogs by using F_ST_ sliding window analysis. Using SNPs of Japanese dogs and African dogs (Table S2) in a vcf file, F_ST_ values for all sites were computed by vcftools v0.1.16 (Danecek, et al. 2011). The average F_ST_ values in windows of 50 SNPs with a 25 SNPs slide across chromosomes were calculated by RStudio version (1.4.1106) (RStudio Team 2020. RStudio: Integrated Development for R. RStudio, PBC, Boston, MA). The Japanese wolf-derived genomic regions were calculated by *f*_dM_ values in windows of 50 SNPs with a 25 SNPs slide across chromosomes using Dsuite (Malinsky, et al. 2021). Then, we extracted regions of overlap between high F_ST_ regions (top 1%) and the Japanese wolf-derived genomic regions (top 1% of *f*_dM_ values).

## Availability of data

The nucleotide sequences were deposited in the DDBJ Sequenced Read Archive.

## Author contributions

JG: data analyses and manuscript writing; NA: next generation sequencing and vcf file preparation of modern samples; XX: assessing DNA damage patterns; YM: genomic DNA extraction of Shiba individuals and next generation sequencing; SM: preparation of an alignment of mitochondrial DNA sequences and manuscript editing; HT: archeological information for discussion and manuscript editing; NI preparation of genomic DNA of the Japanese wolves and Japanese dogs and manuscript editing; YT: research concept, research plan, next generation sequencing, vcf file preparation, data analyses, and manuscript writing.

---

**Figure S1.**
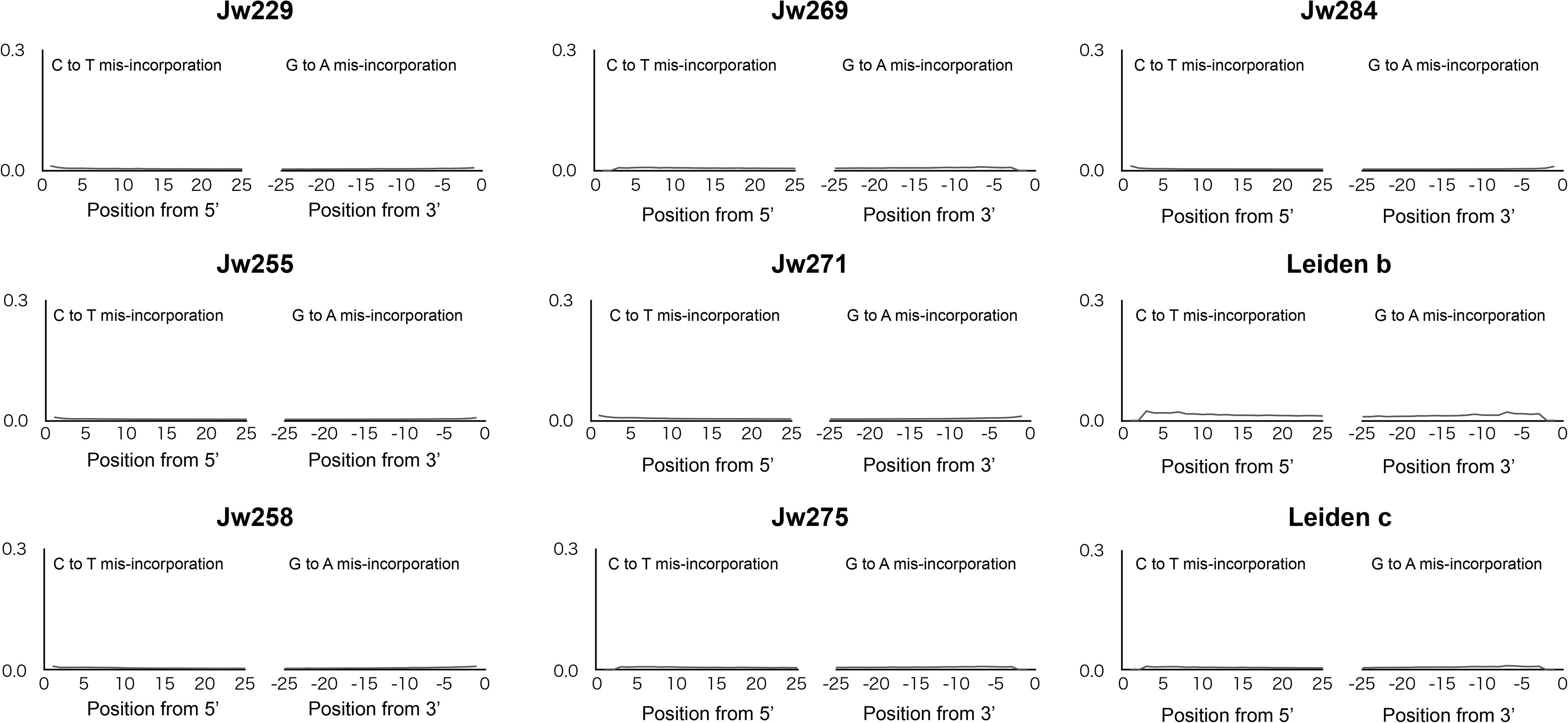
C to T and G to A frequency of mis-incorporation at 3’ and 5’ end of reads determined in this study.

**Figure S2.**
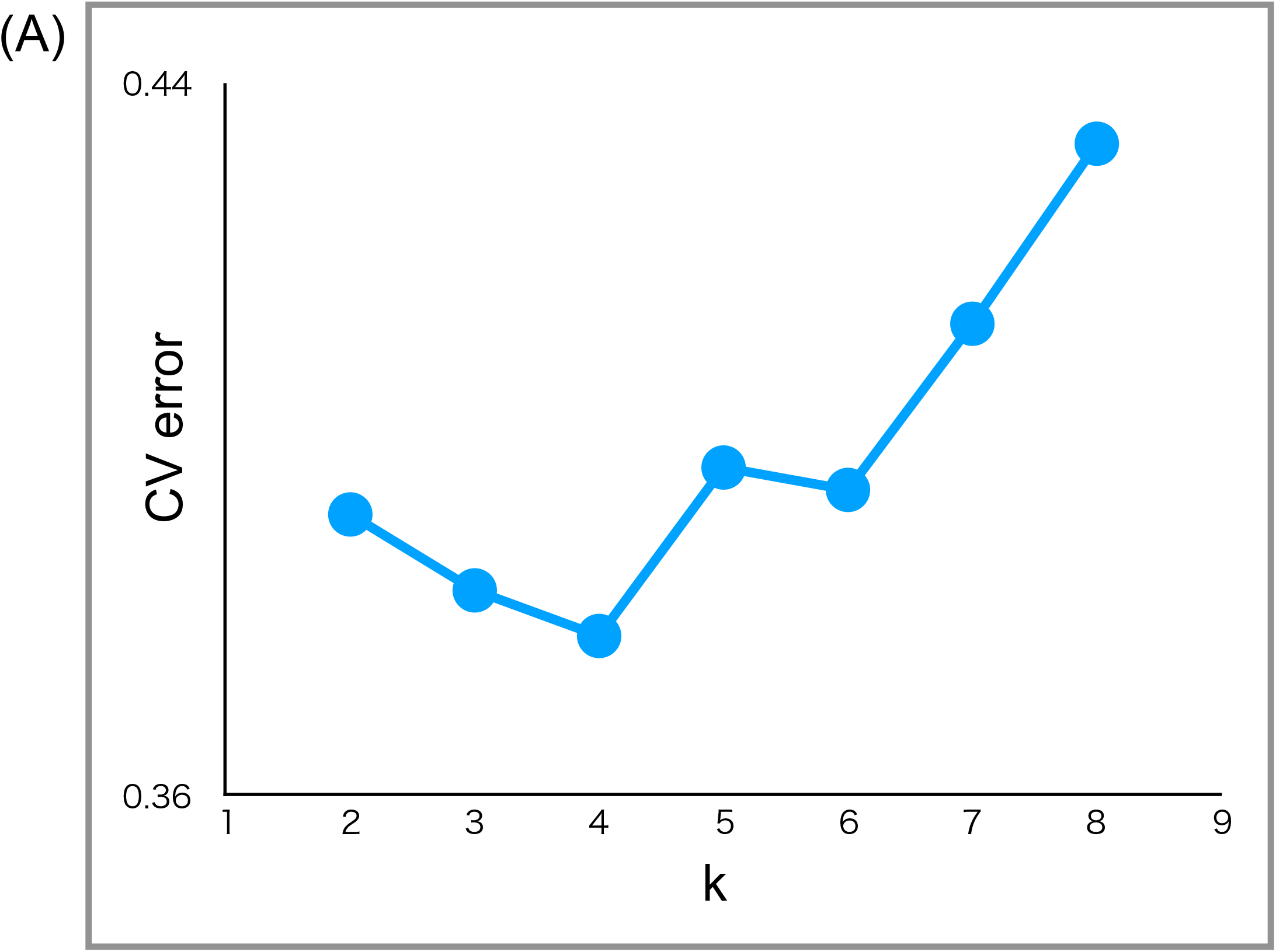
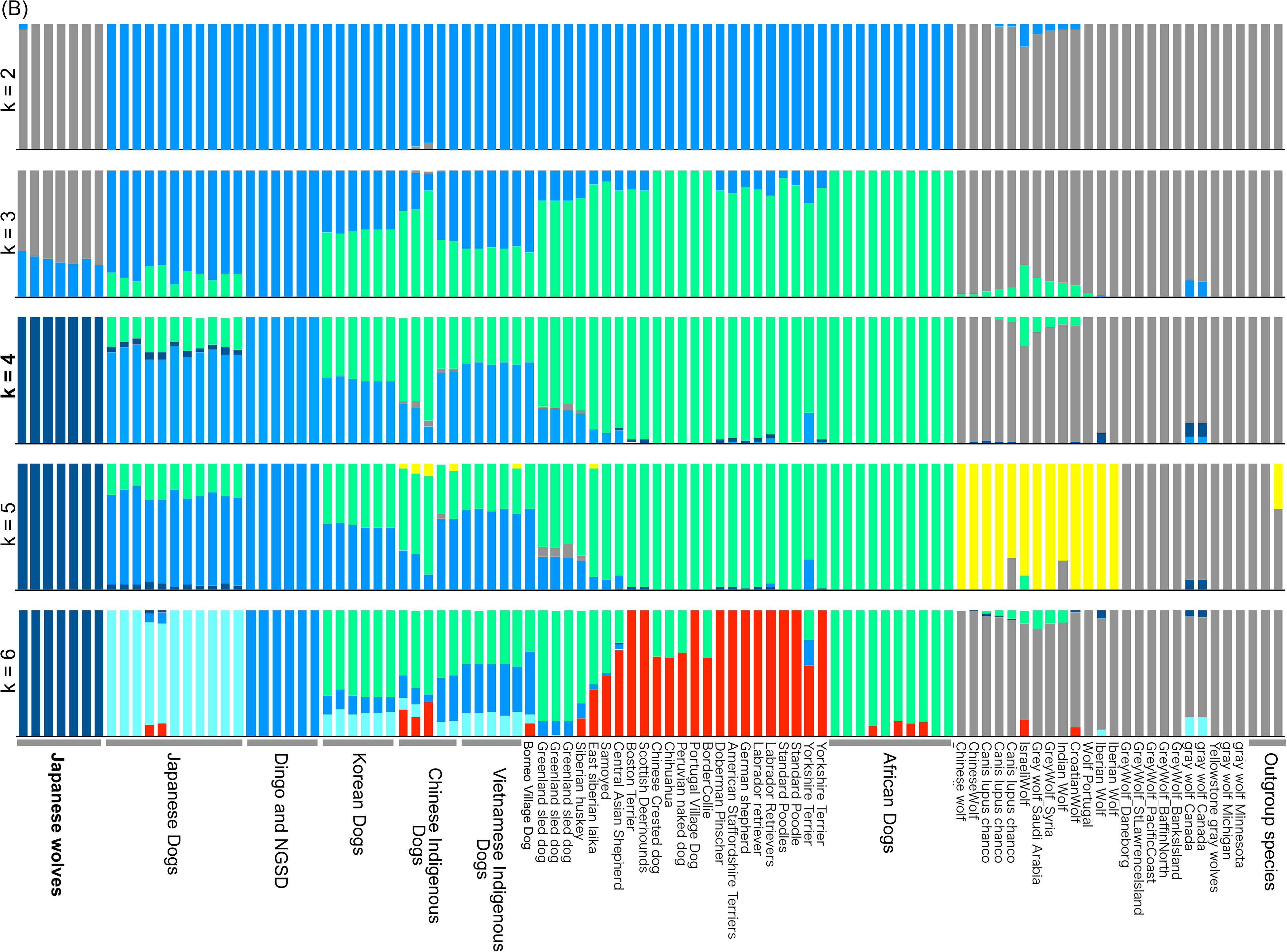
(A) Cross validation (CV) values for ADMIXTURE analysis of SNP data. (B) ADMIXTURE results based on SNP data for K = 2-6 (see Table S2 for sample information).

**Figure S3.**
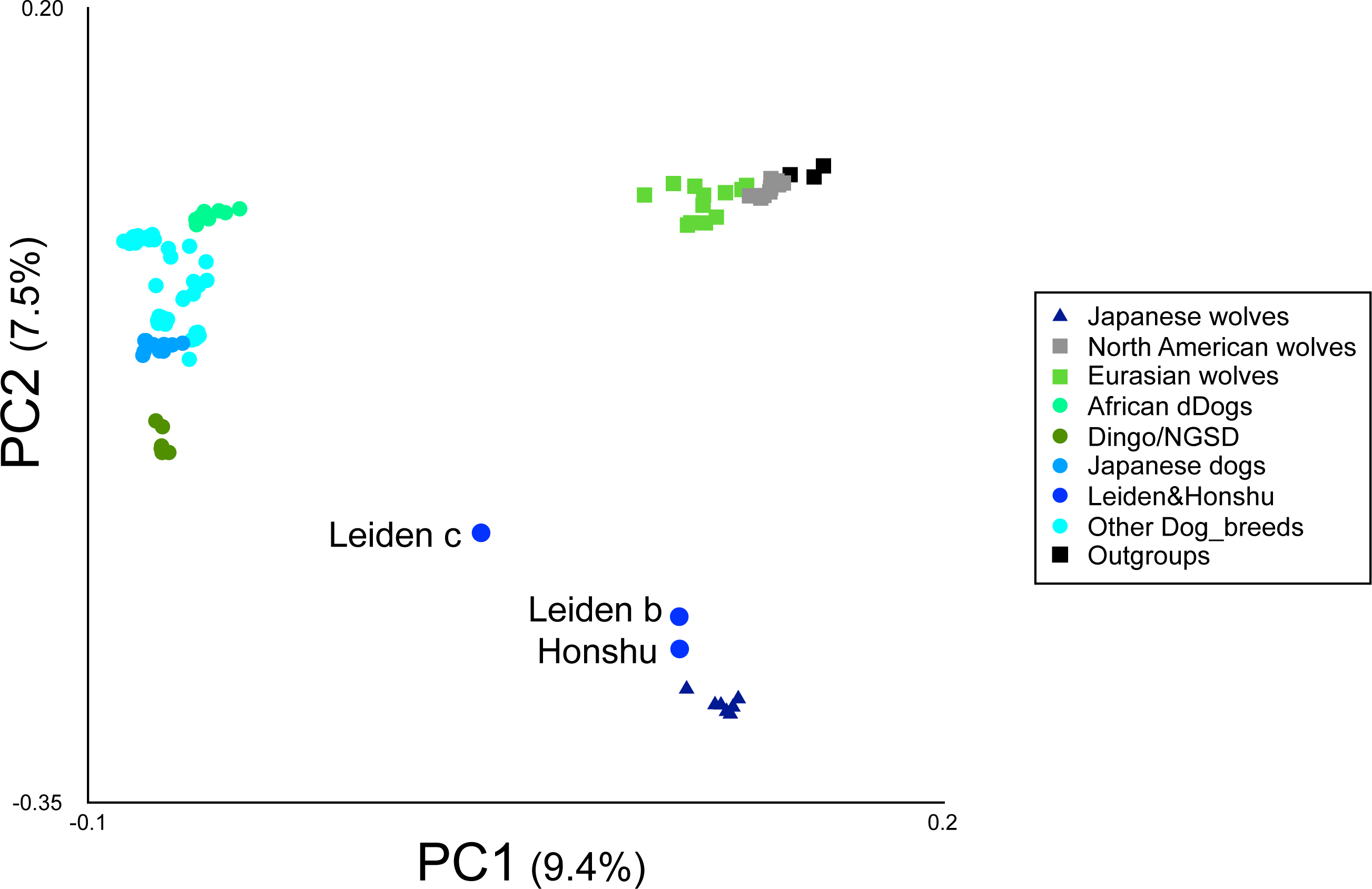
Principal Components Analysis (PC1 versus PC2) of 103 samples based on 103,432 SNPs (see Table S2 for sample information). Colored circle, square, and triangle correspond to the names of dogs or wolves in the panel.

**Figure S4.**
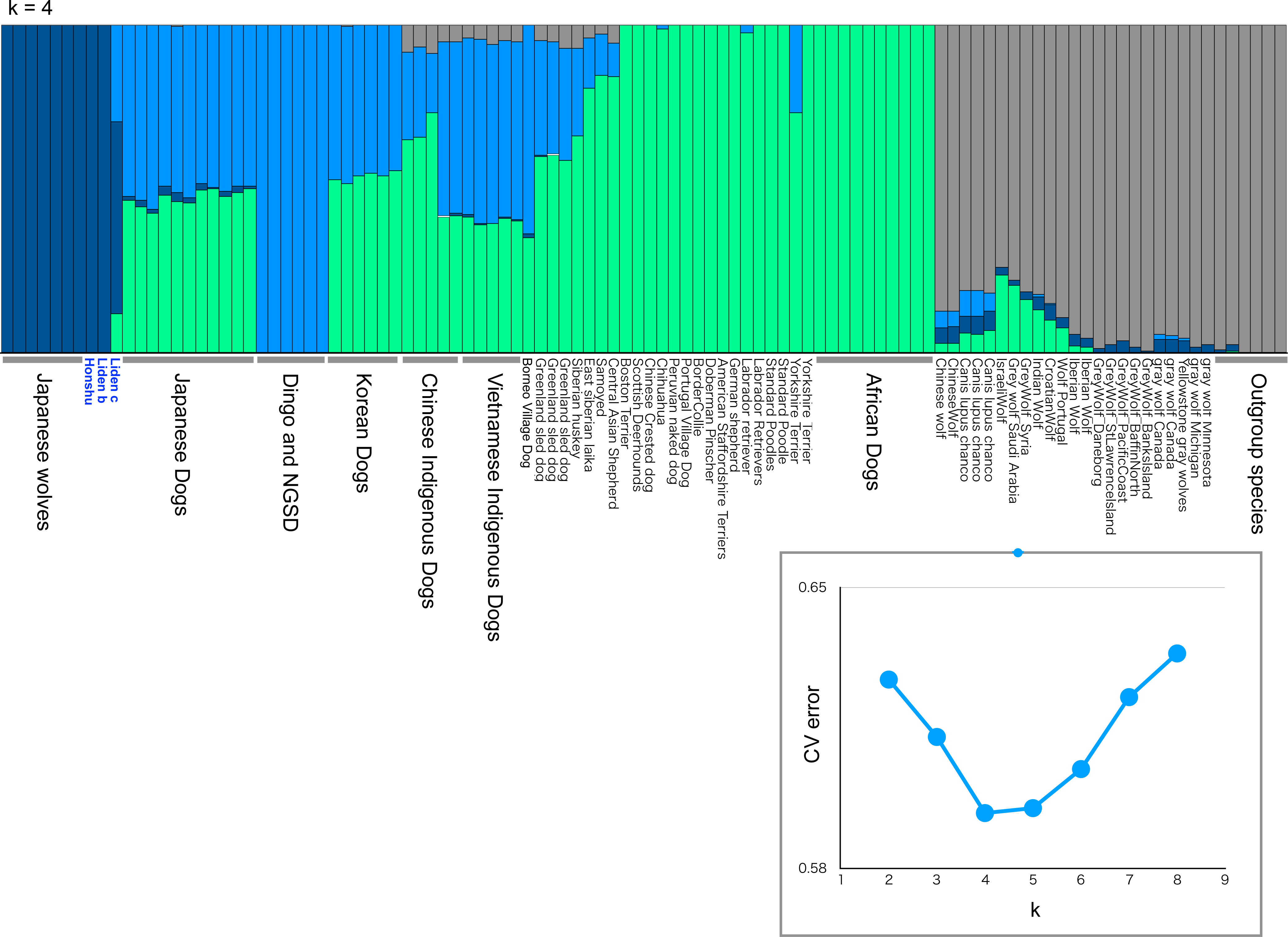
An ADMIXTURE result based on SNP data for K = 4 (see Table S2 for sample information). Cross validation (CV) values for ADMIXTURE analysis of SNP data is shown in the panel.

**Figure S5.**
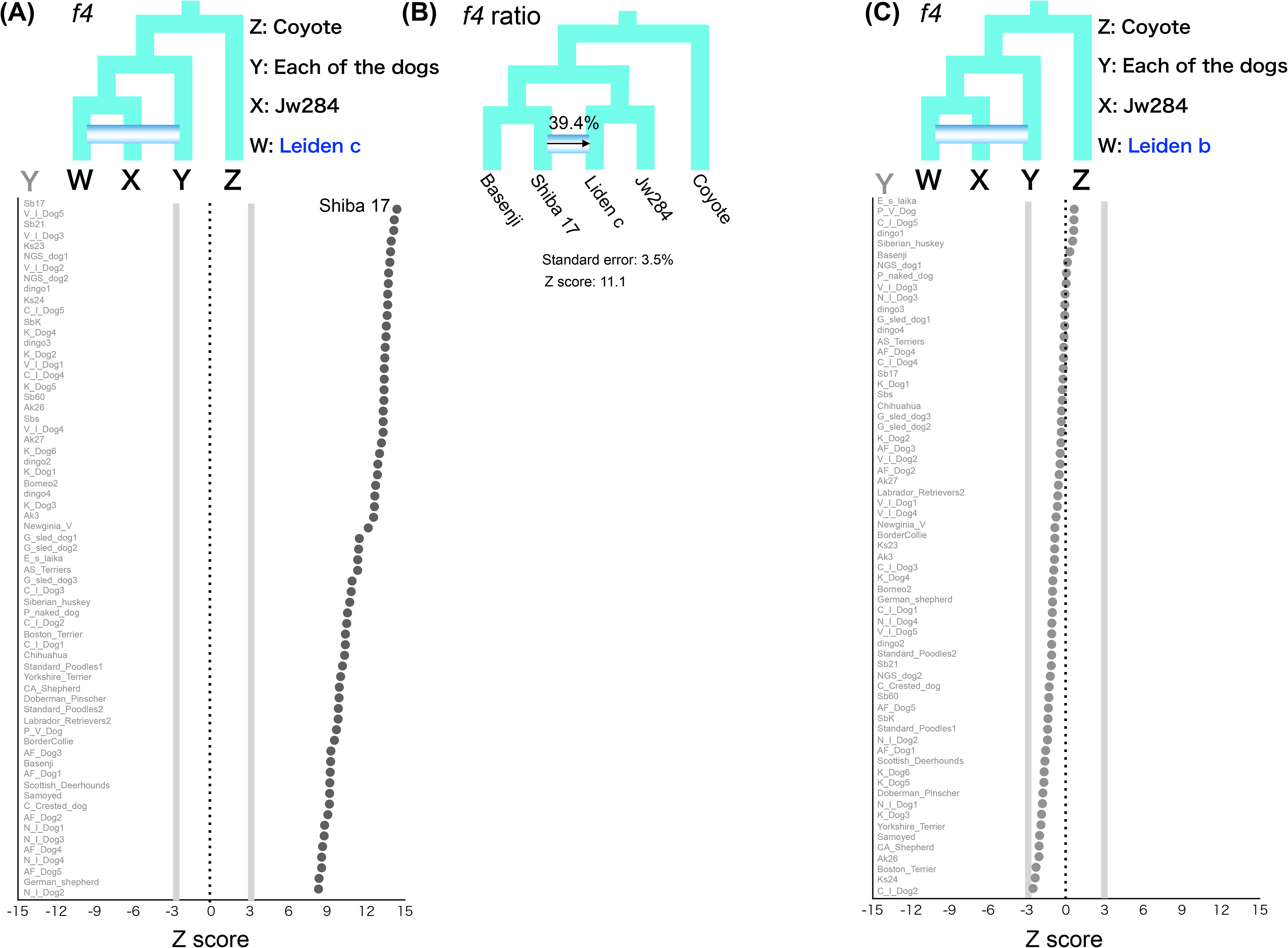
(A) *f4*-statistics testing the relationships between the Japanese wolf (Jw284), Leiden c, and all other dogs. Each Z score is plotted in order of highest to lowest value from the top, and the names of dogs are shown on the left side of each panel. Gray lines show the Z score −3 and 3. Leiden c shows genetic affinity with all other dogs (Z score > 3). (B) *f4*-ratio test to estimate proportion of genome introgression from a Japanese dog (Shiba 17) to Leiden C. (C) *f4*-statistics testing the relationships between the Japanese wolf (Jw284), Leiden b, and all other dogs. The genetic affinities of Leiden b with all other dogs are rejected (−3 < Z score < 3).

**Figure S6.**
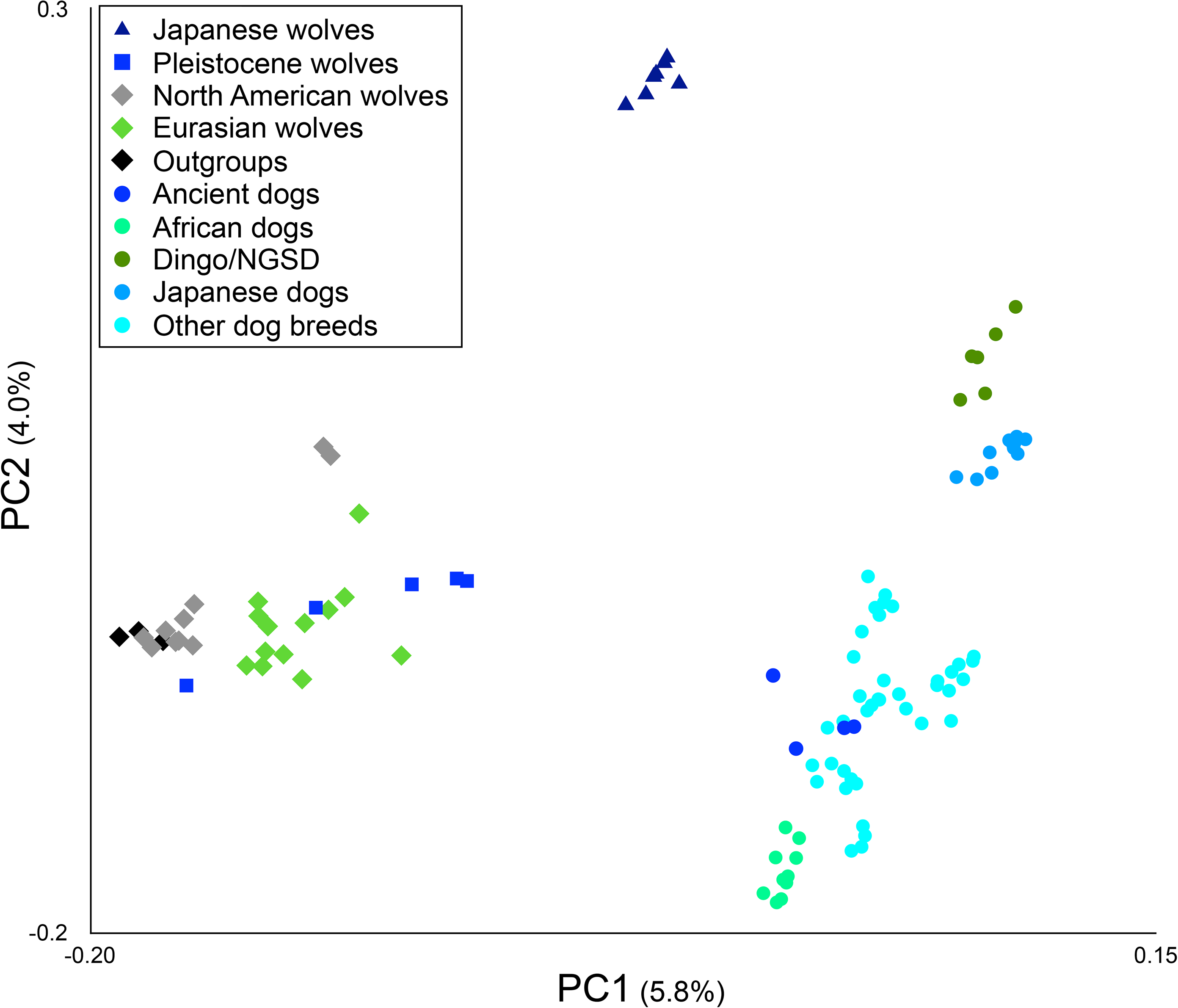
Principal Components Analysis (PC1 versus PC2) of 109 samples based on 100,588 SNPs (transversion sites, see Table S2 for sample information). Colored circle, square, and triangle correspond to the names of dogs or wolves in the panel.

**Figure S7.**
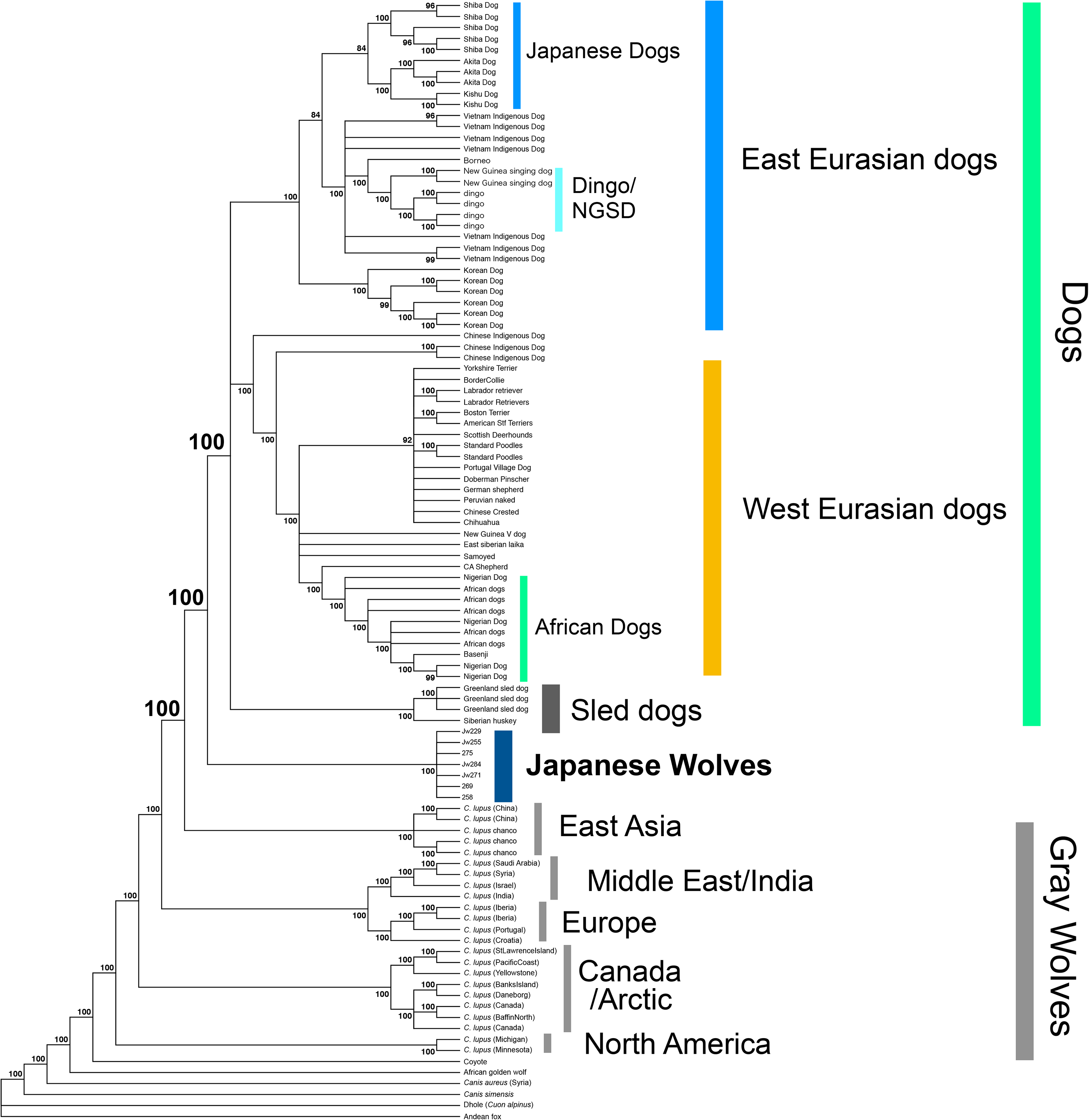
Maximum likelihood tree based on 489,524 SNPs. Node labels indicate bootstrap replicates.

**Figure S8.**
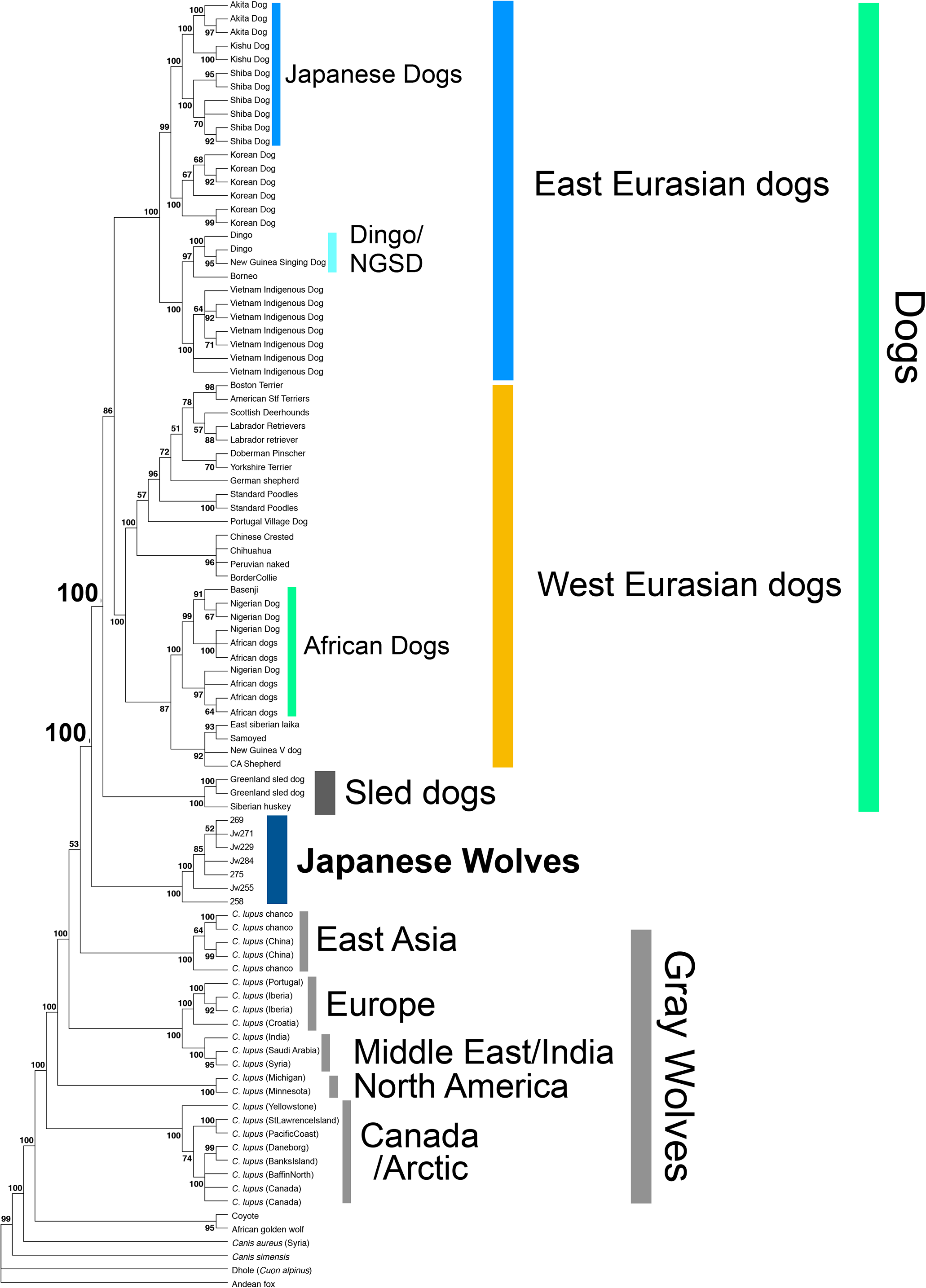
Phylogenetic tree constructed by SVDquartets based on 489,524 SNPs. Node labels indicate bootstrap replicates.

**Figure S9.**
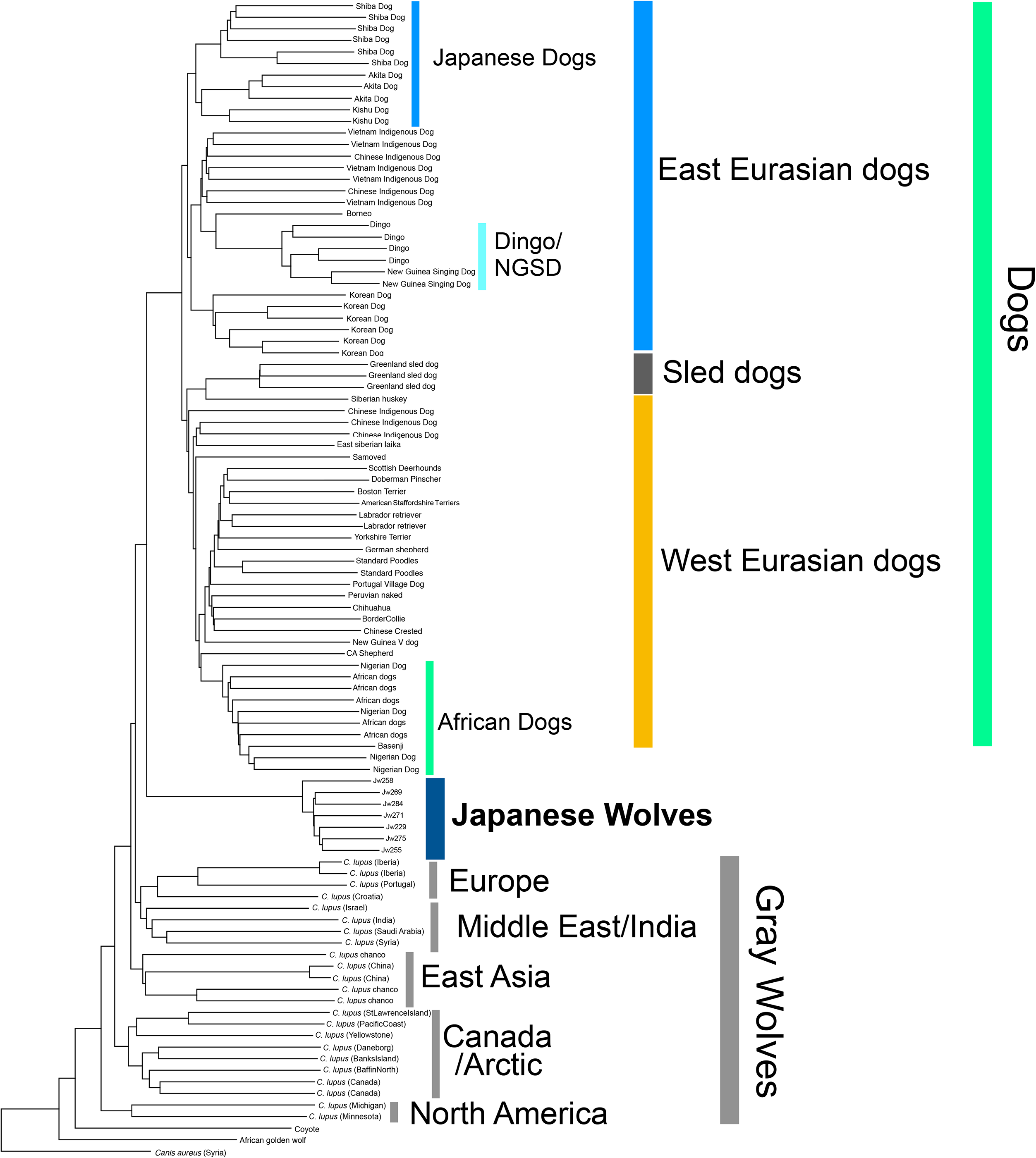
NJ tree based on IBS matrix from 1,992,260 SNPs.

**Figure S10.**
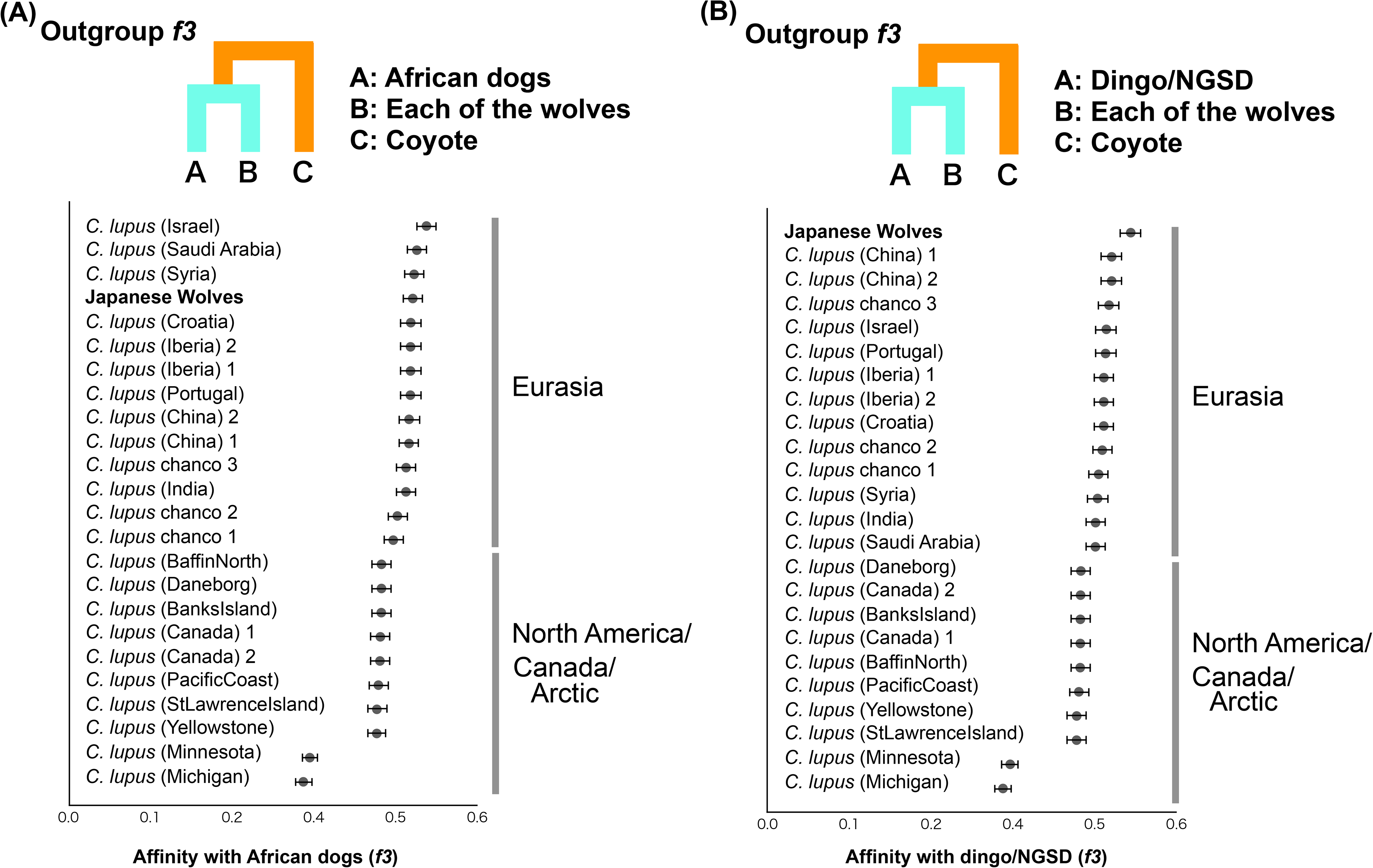
Shared genetic drift between African dogs (A) and Dingo/NGSD (B) and gray wolves measured by outgroup *f*3 statistics. Each of the African dogs and Dingo/NGSD individuals were used as populations. Each *f3* statistical value is plotted in order of highest to lowest value from the top, and the names of the wolves are shown on the left side of each panel. Error bars represent standard errors.

**Figure S11.**
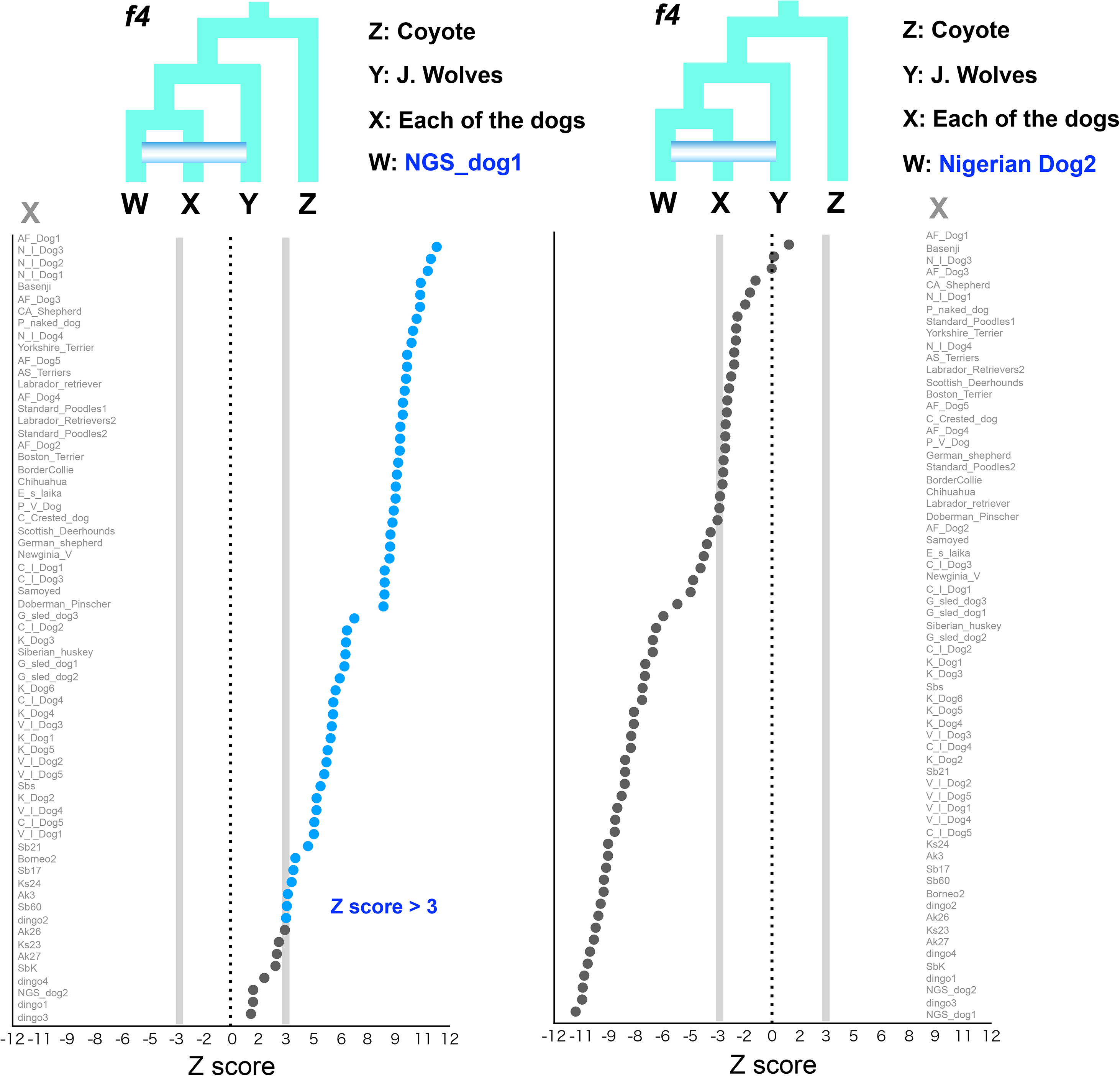
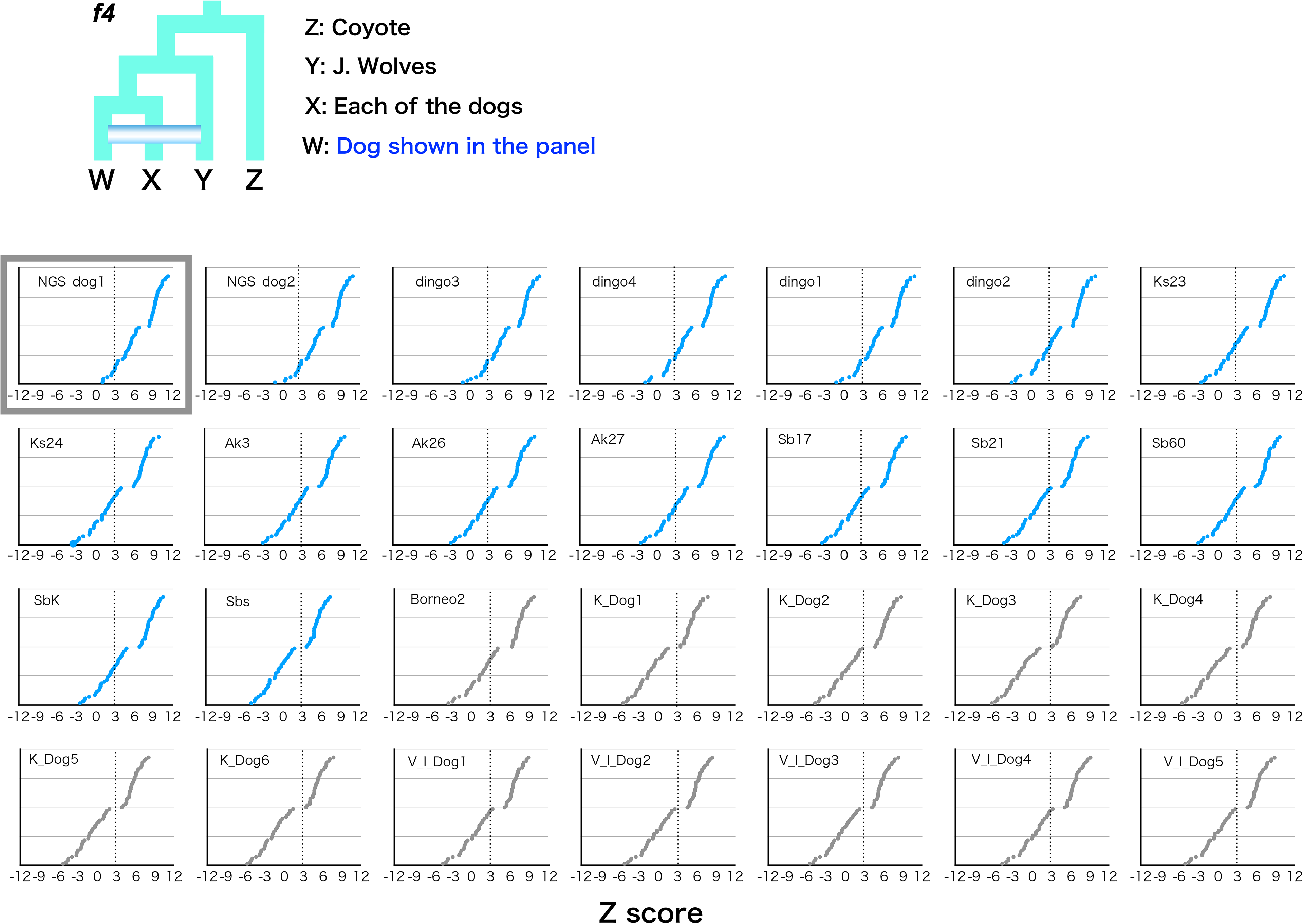
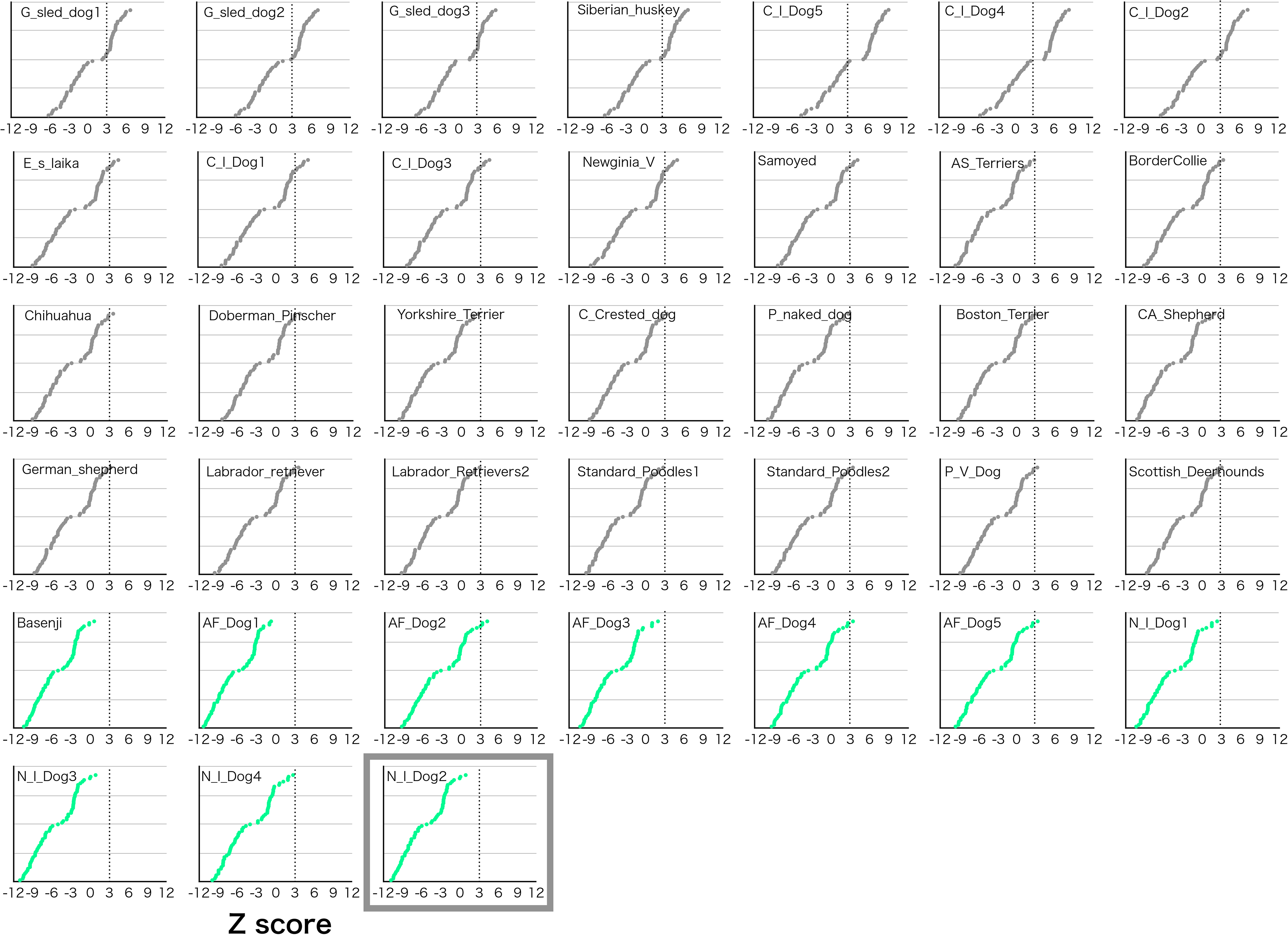
*f4* statistics testing the genetic affinity of the Japanese wolf with all other dogs. All Japanese wolves were used as a population. Z scores for each combination are plotted. We computed *f4* statistics where W in the schematic representation is shown and fixed in each panel and X represented any possible other dogs. Each Z score is listed in order of highest to lowest value from the top. Dotted line shows the Z score 3. Large size graphs of highest (NGS_dog1) and lowest (Nigerian Dog2) affinity to the Japanese wolf are show in the first two panels. The names of the dogs are shown on the left or right sides of the panels. Small size panels of these two are surrounded by gray squares.

**Figure S12.**
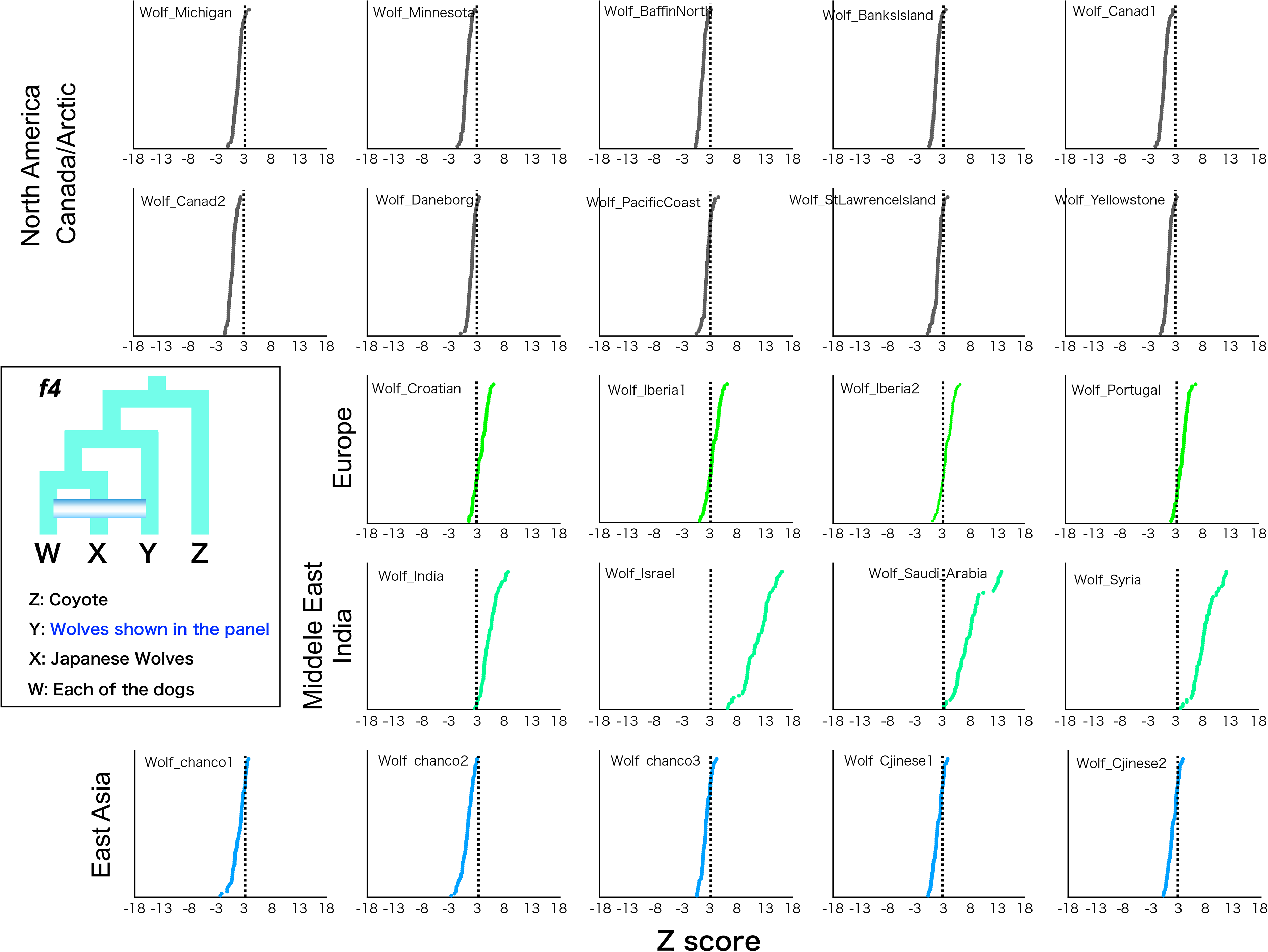
*f4* statistics testing the genetic affinity of gray wolves shown in each panels with all dogs. Z scores for each combination are plotted in order of highest to lowest value from the top. We computed *f*4 statistics where Y in the schematic representation is shown and fixed in each panel and W represented any possible other dogs. Dotted line shows the Z score 3.

**Figure S13.**
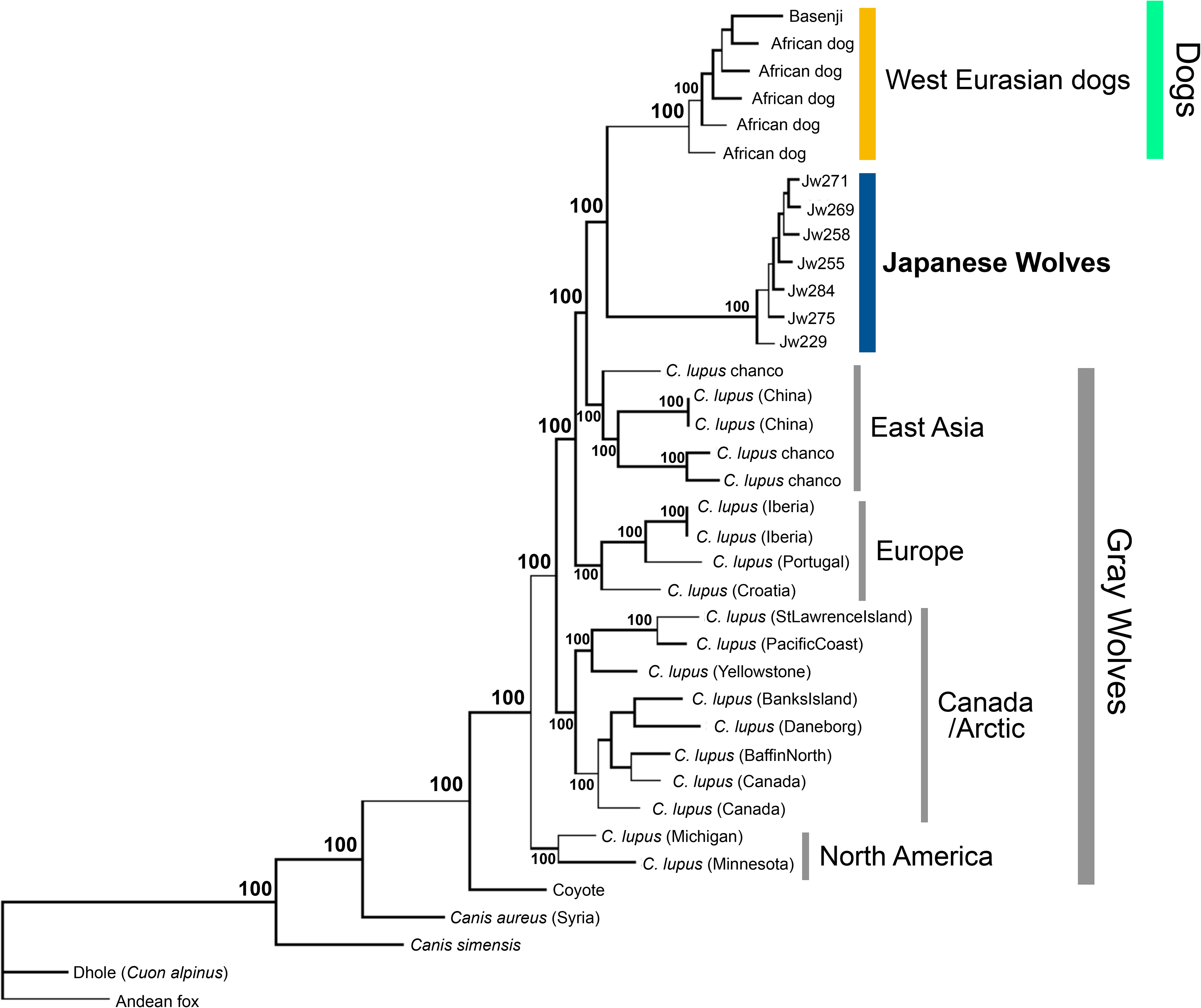
Maximum likelihood tree based on 2,065,200 SNPs. Node labels indicate bootstrap replicates.

**Figure S14.**
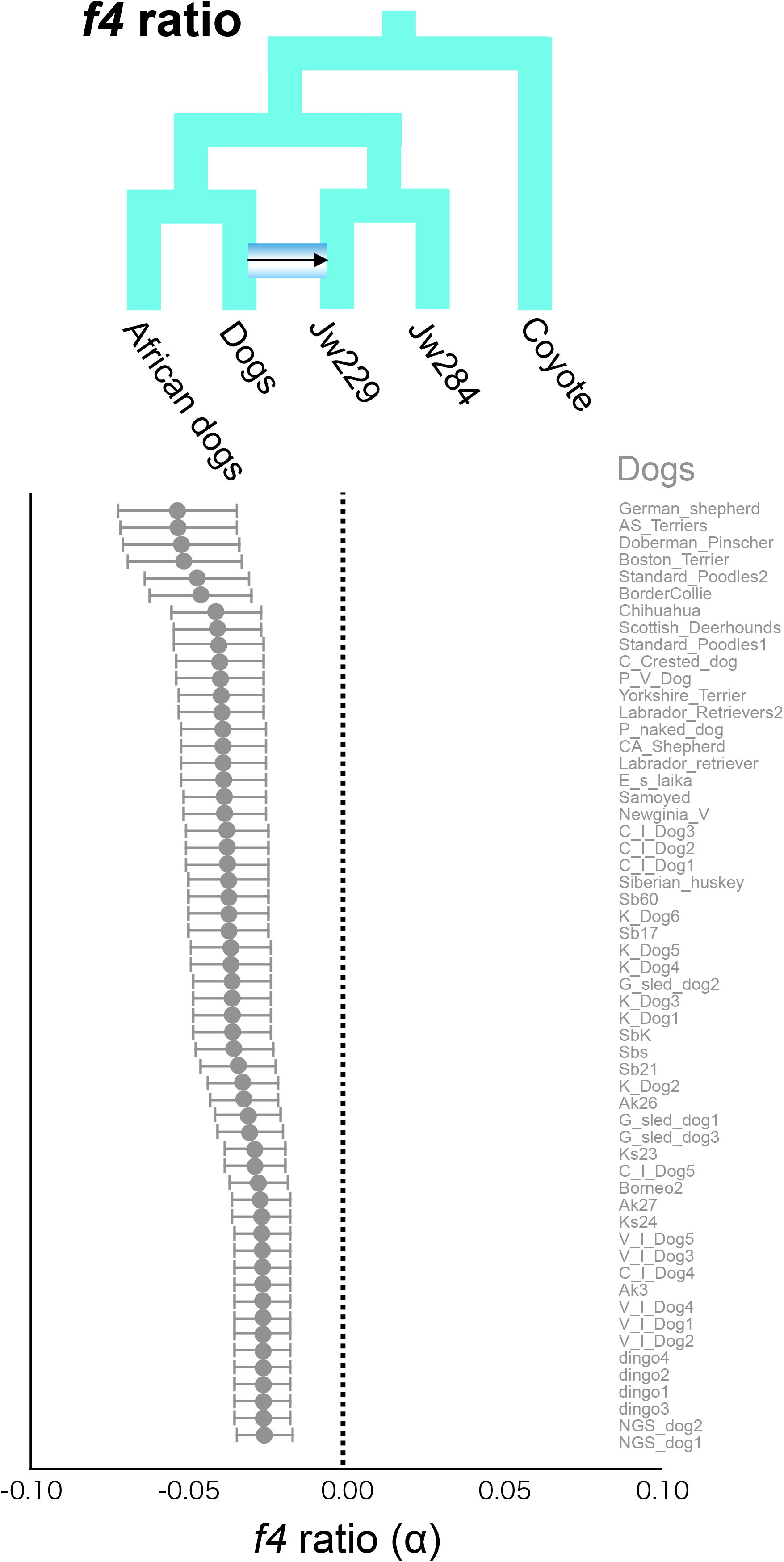
*f4*-ratio test to estimate proportion of genome introgression from dogs to the Japanese wolf. Each *f4*-ratio α value is plotted in order of lowest to highest value from the top, and the names of the dogs are shown on the right side of the panel. Error bars represent standard errors.

**Figure S15.**
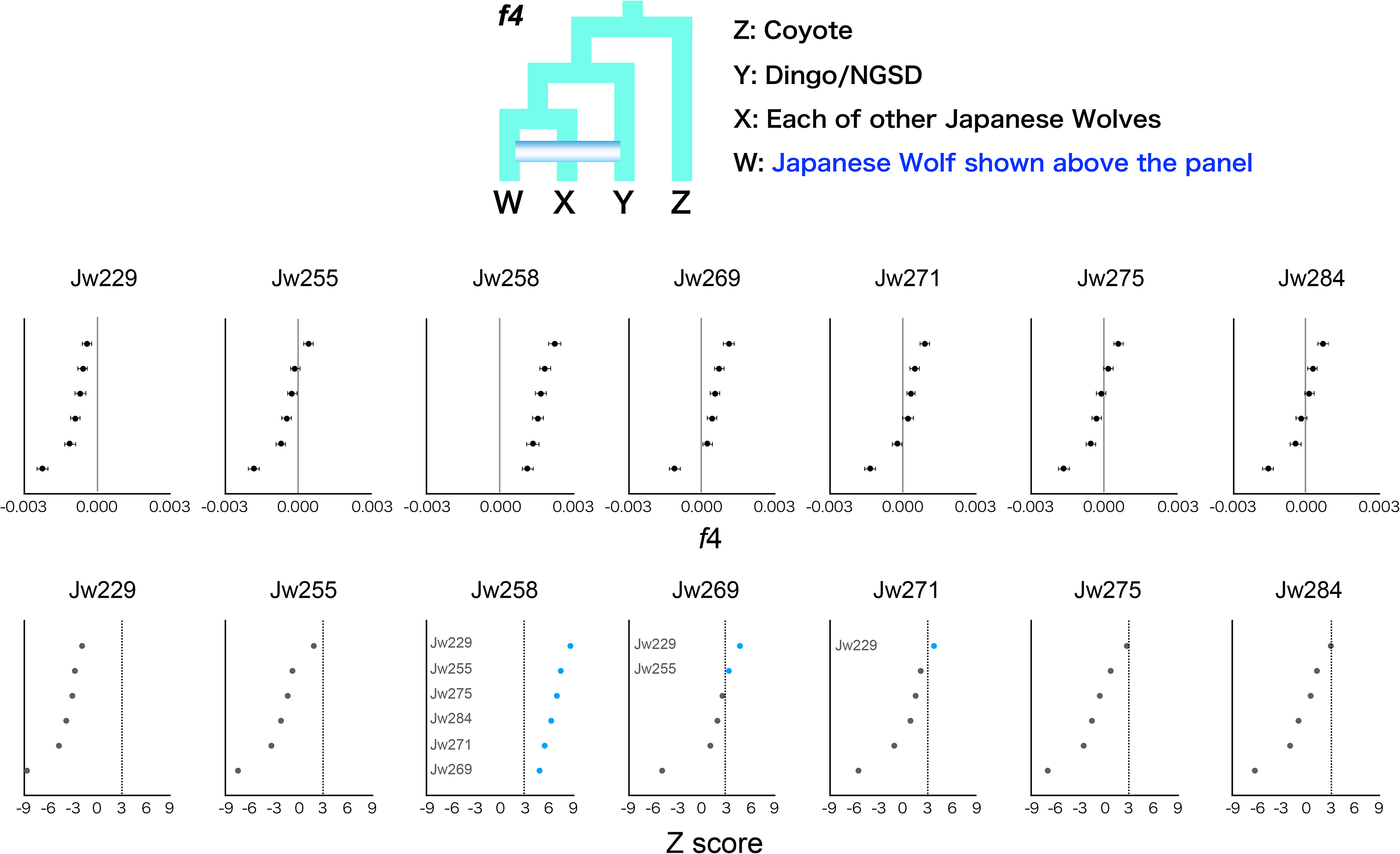
(A) *f4* statistics testing the difference of the affinity to dingo/NGSD between the Japanese wolf individuals. *f4* statistics (upper panels) and Z score (lower panels) value is plotted in order of highest to lowest value from the top. Z score above 3 is colored in blue. When the Japanese wolf individual showing a significant affinity to dingo/NGSD, the names of the Japanese wolf individual at the position X in the schematic representation are shown on the left side of Z score panel.

**Figure S16.**
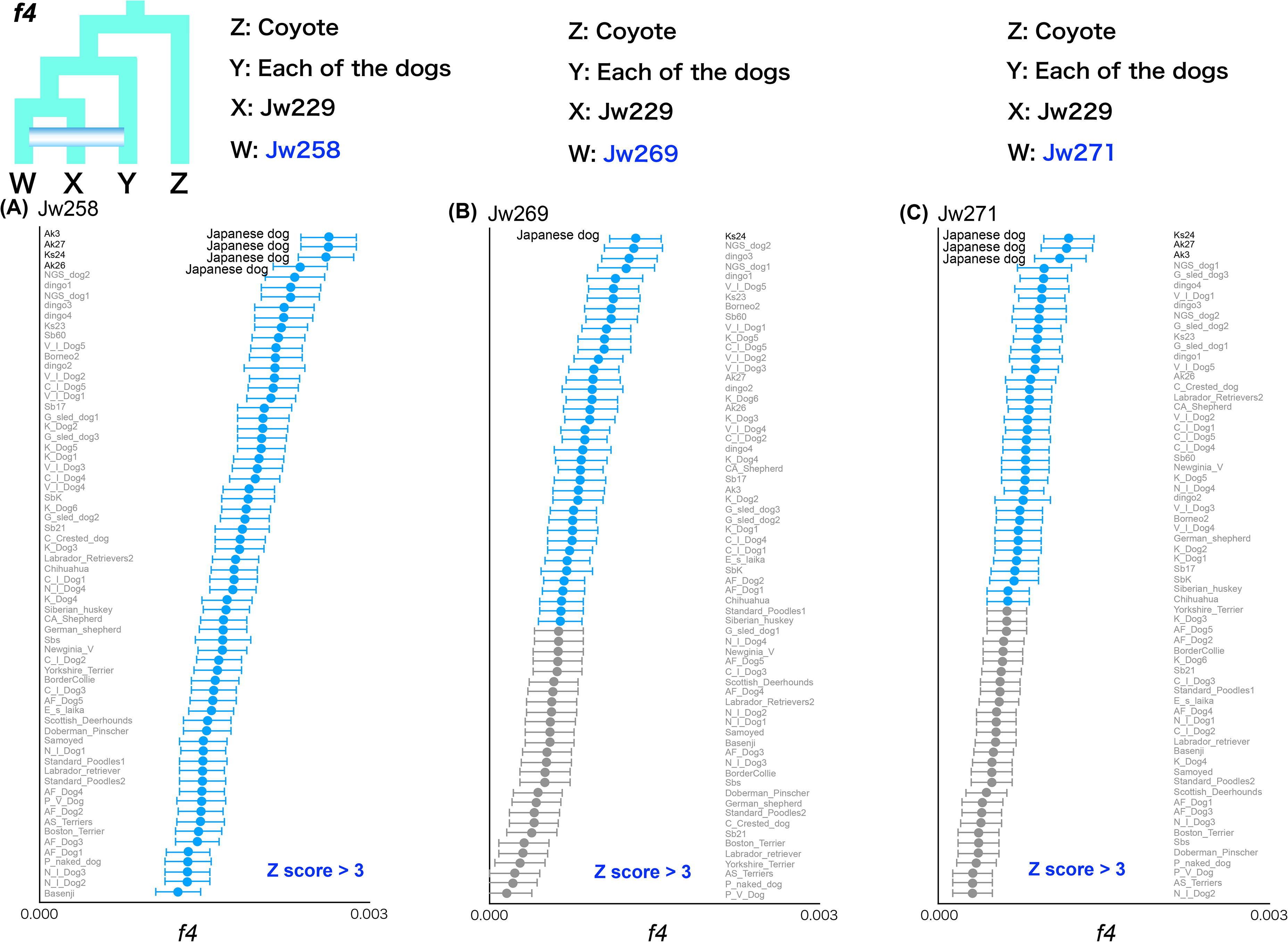
*f4* statistics testing the relationships between (A) Jw258, (B) Jw269, and (C) Jw271 and all dog individuals. Each Z score is plotted in order of highest to lowest value from the top, and the names of dogs are shown on theft or right sides of each panel (see Table S2). Z score above 3 is colored in blue.

**Figure S17.**
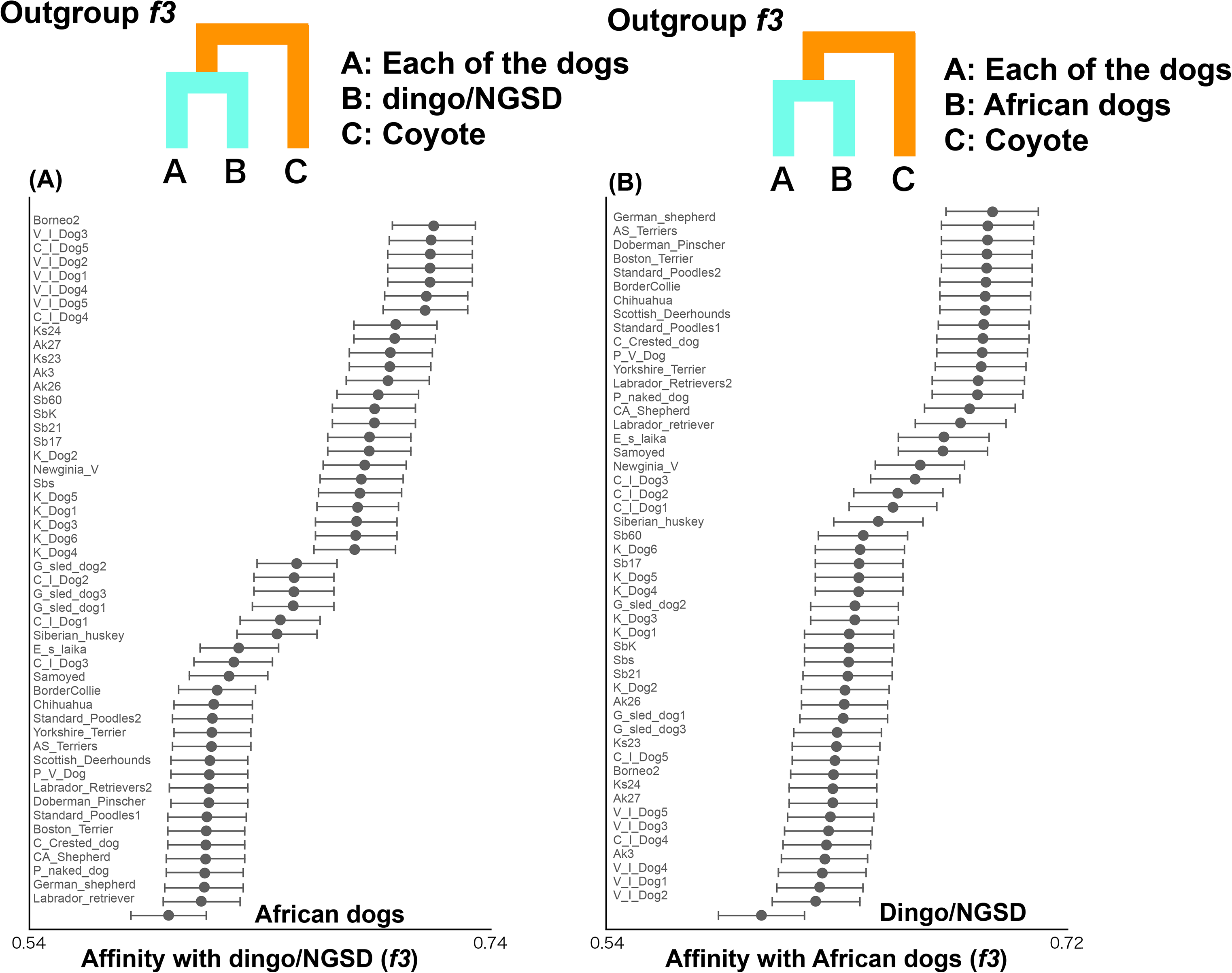
Shared genetic drift between (A) dingo/NGSD and (B) African dogs and all dogs measured by outgroup *f*3 statistics. Each of the African dogs and dingo/NGSD individuals were used as populations. Each *f3* value is plotted in order of highest to lowest value from the top, and the names of the dogs are shown on the left side of the panel. Error bars represent standard errors.

**Figure S18.**
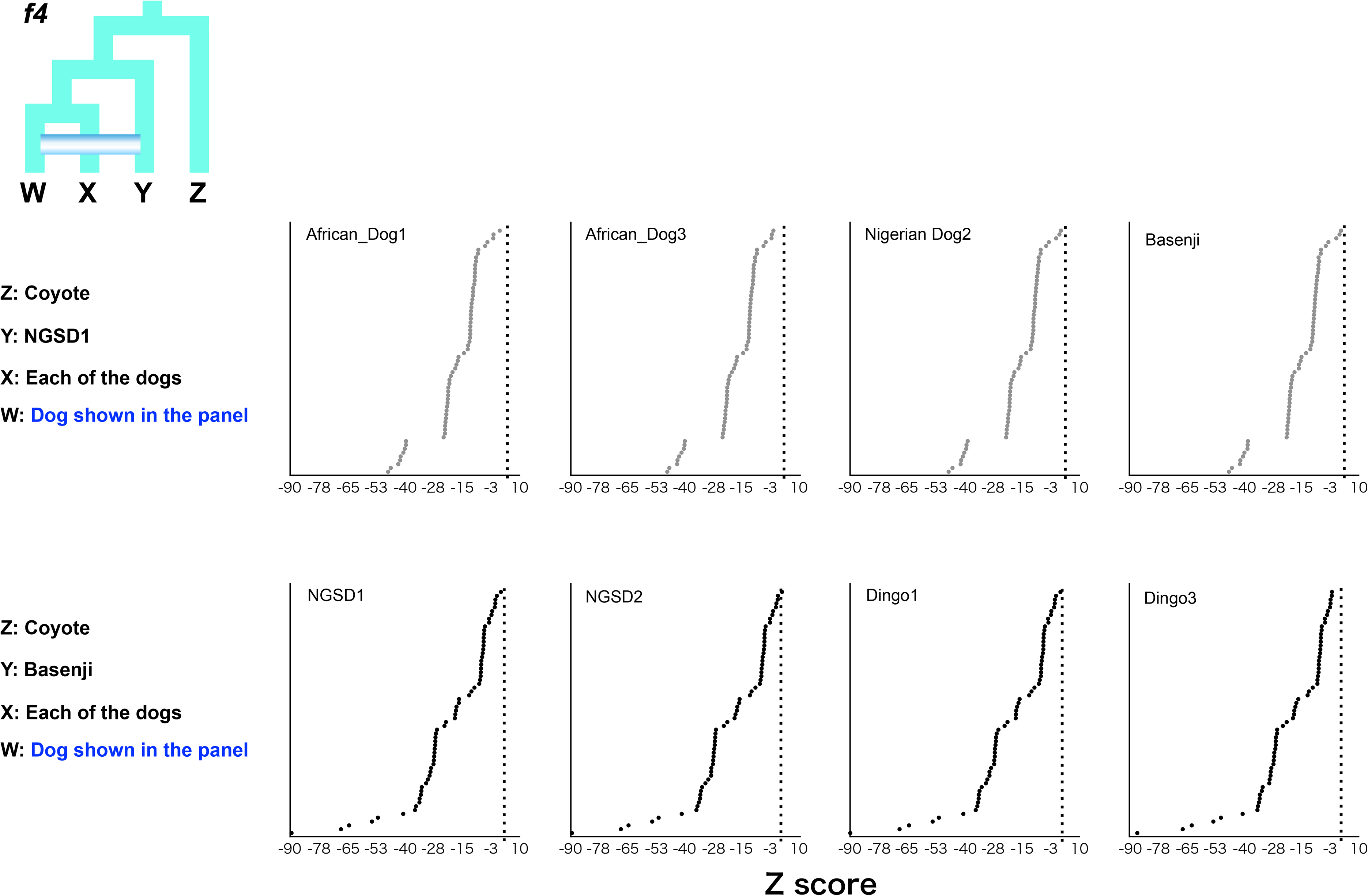
*f4* statistics testing the affinity of NGSD1 with African dogs (upper panels) and that of Basenji with dingo/NGSD dogs (lower panels). All Z scores were under 3.

**Figure S19.**
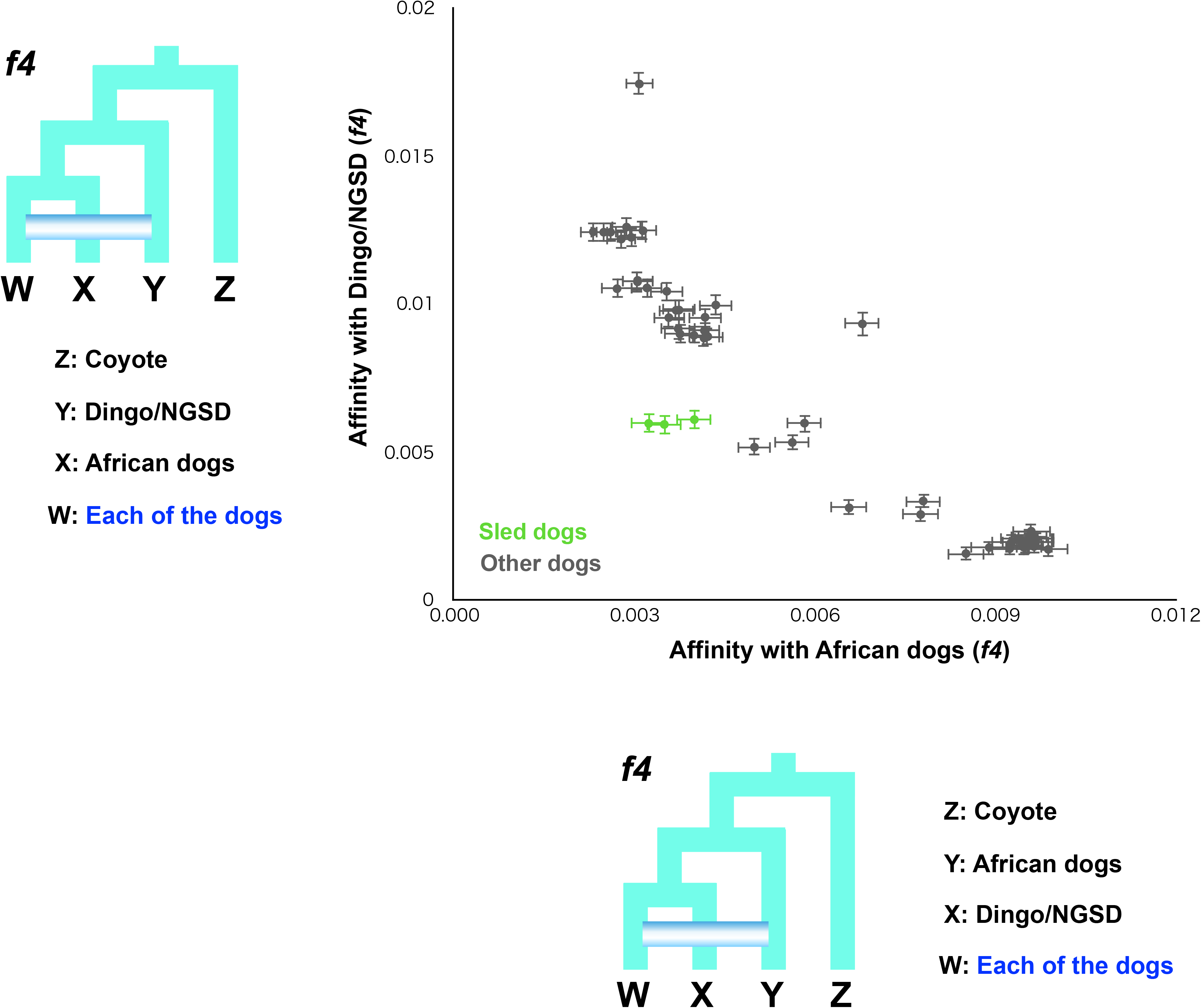
*f4* statistics testing whether dogs share more alleles with African dogs (x-axis) or dingo/NGSD (y-axis) compared with dingo/NGSD and African dogs, respectively. Dots show the *f4* statistics, and horizontal and vertical error bars represent standard errors for the test with the African dogs (x-axis) and dingo/NGSD (y-axis), respectively. Each of the African dogs and dingo/NGSD individuals were used as populations.

**Figure S20.**
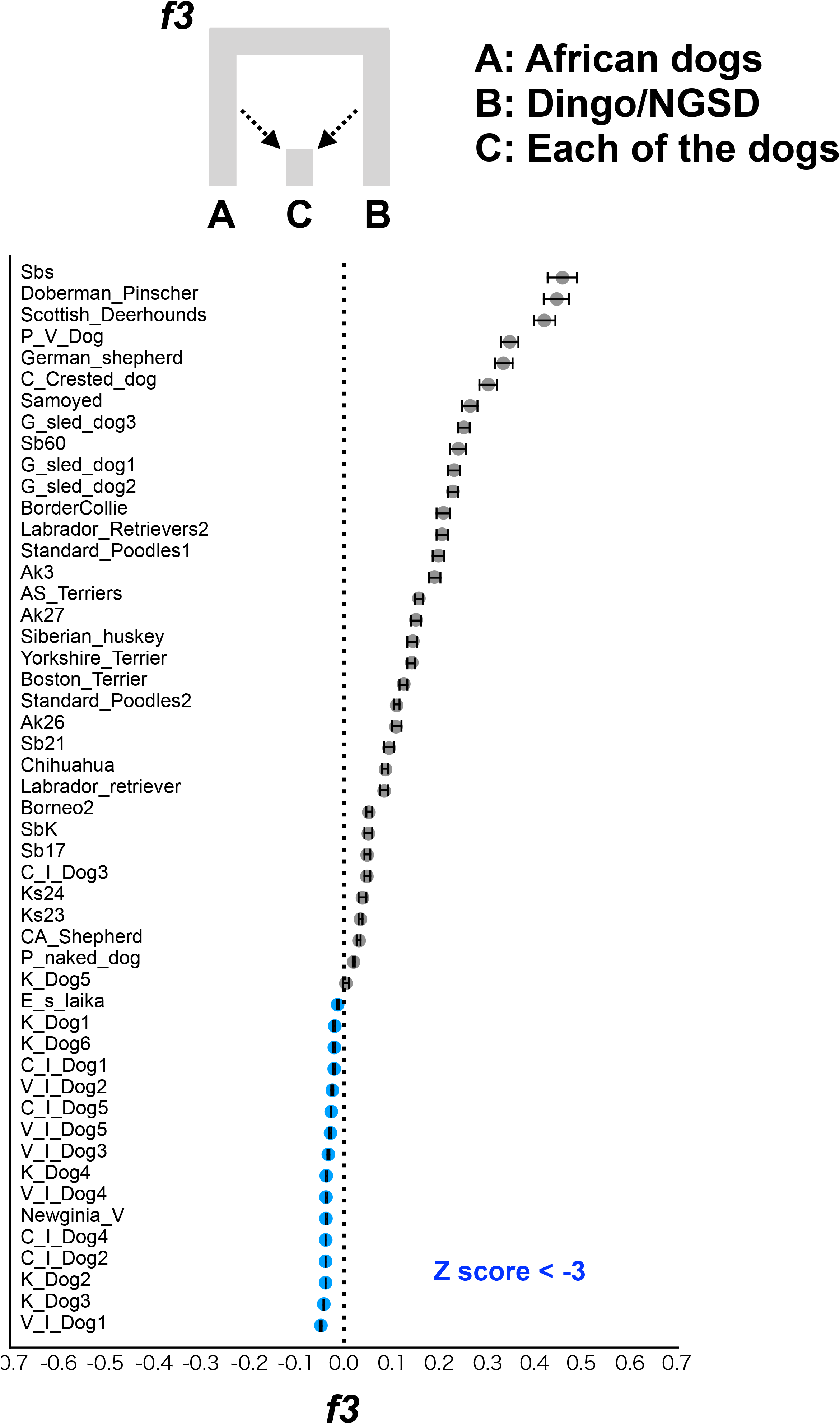
*f3* statistics testing the genomic mixture of African and Dingo/NGSD dogs in all dogs. Z score under −3 is colored in blue. Each of the African dogs and Dingo/NGSD individuals were used as populations. Each *f3* value is plotted in order of highest to lowest value from the top, and the names of the dogs are shown on the left side of the panel. Error bars represent standard errors.

**Figure S21.**
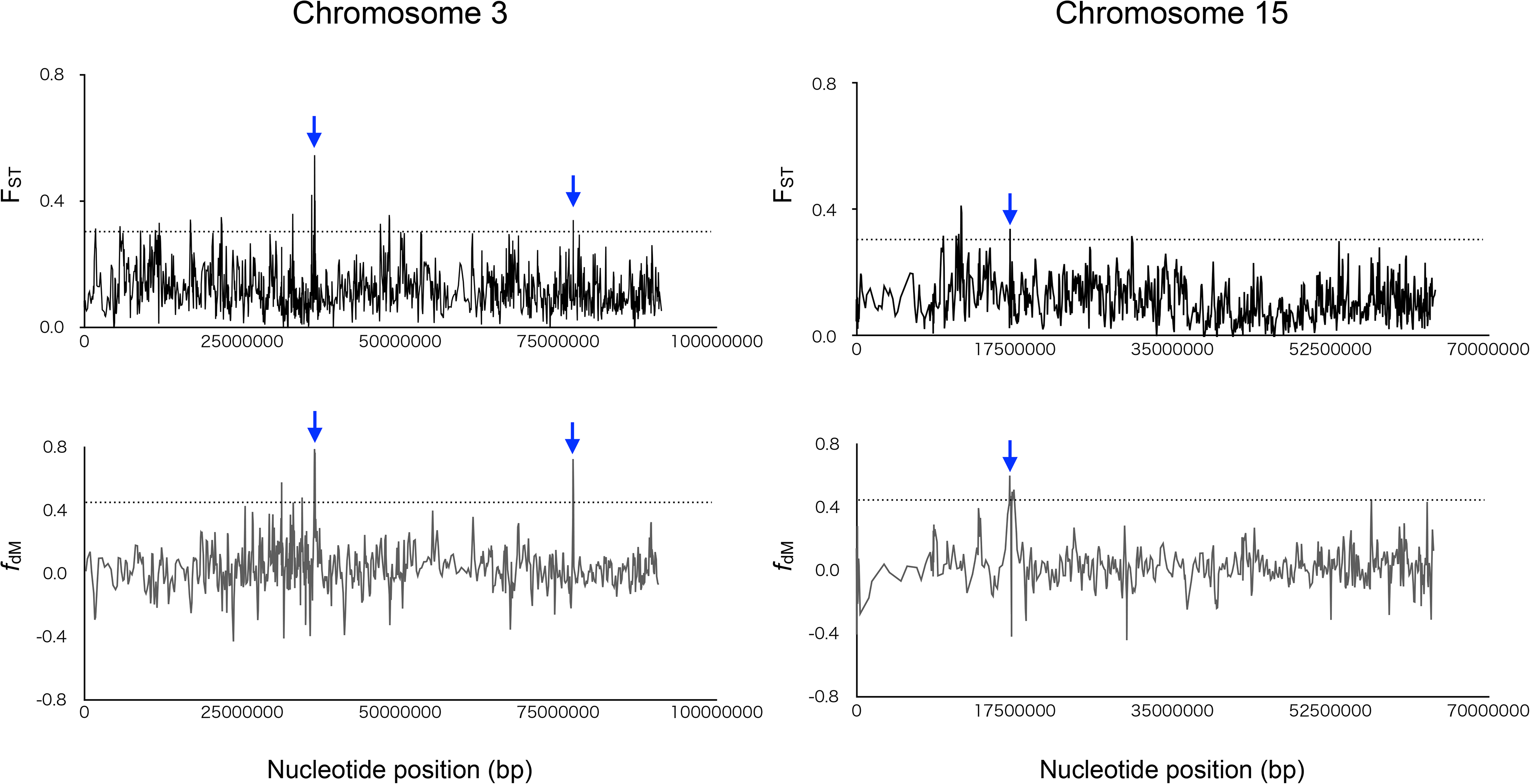
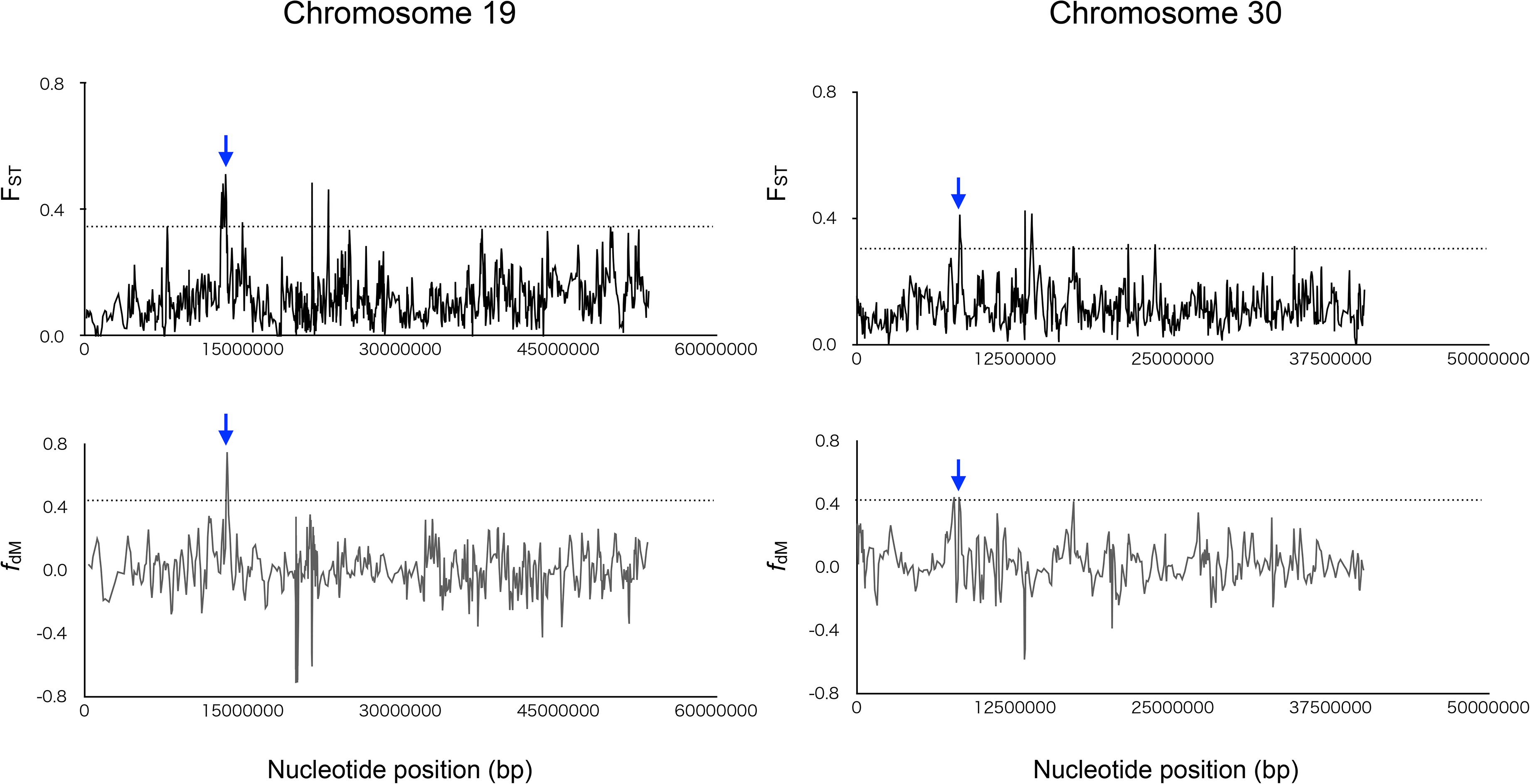
Sliding window analyses of the F_ST_ values (y-axis: upper panel) and *f*_dM_ (y-axis: lower panel) in windows of 50 SNPs using a 25 SNPs slide across scaffolds (x-axis). Dashed lines show the 99th percentiles. Blue arrow indicate overlapping regions above 99th percentiles between upper and lower panels.

**Table S1.** Determined sequences in this study

| Wolf/Dog | ID | Period | Isolation site (Prefecture) | Mapped reads count | Total mapped reads |
| --- | --- | --- | --- | --- | --- |
| Japanese Wolves | Leiden_b (Jentink 1887 b RMNH.MAM.39183) | Edo | Leiden | 149 M | 22.35 Gb |
| Japanese Wolves | Leiden_c (Jentink 1887 c RMNH.MAM.39181) | Edo | Leiden | 188 M | 28.2 Gb |
| Japanese Wolves | Jw255 | Edo-Meiji | Yamanashi Pref. | 1881 M | 282.2 Gb |
| Japanese Wolves | Jw271 | Edo-Meiji | Iwate Pref. | 820 M | 123 Gb |
| Japanese Wolves | Jw284 (ZMB_Mam_048817) | Meiji | Berlin | 1237 M | 185.6 Gb |
| Japanese Wolves | Jw275 | Edo-Meiji | Shimane Pref. | 134 M | 46.5 Gb |
| Japanese Wolves | Jw229 | Edo | Kochi Pref. | 661 M | 99.2 Gb |
| Japanese Wolves | Jw258 | Edo-Meiji | Nagano Pref. | 262 M | 110 Gb |
| Japanese Wolves | Jw269 | Edo-Meiji | Nagano Pref. | 193 M | 61.8 Gb |
| Dog (Akita) | Akita26 | Modern | — | 847 M | 127.1 Gb |
| Dog (Akita) | Akita27 | Modern | — | 808 M | 121.2 Gb |
| Dog (Akita) | Akita3 | Modern | — | 772 M | 115.8 Gb |
| Dog (Kishu) | Kishu23 | Modern | — | 832 M | 124.8 Gb |
| Dog (Kishu) | Kishu24 | Modern | — | 634 M | 95.1 Gb |
| Dog (Shiba) | Shiba17 | Modern | — | 632 M | 94.8 Gb |
| Dog (Shiba) | Shiba21 | Modern | — | 517 M | 77.6 Gb |
| Dog (Shiba) | Shiba60 | Modern | — | 668 M | 100.2 Gb |
| Dog (Shiba) | ShibaKuro | Modern | — | 472 M | 70.8 Gb |
| Dog (Shiba) | ShibaShiro | Modern | — | 697 M | 104.6 Gb |
| Dog (Shiba) | Jm | Modern | — | 391 M | 58 Gb |

**Table S2.**
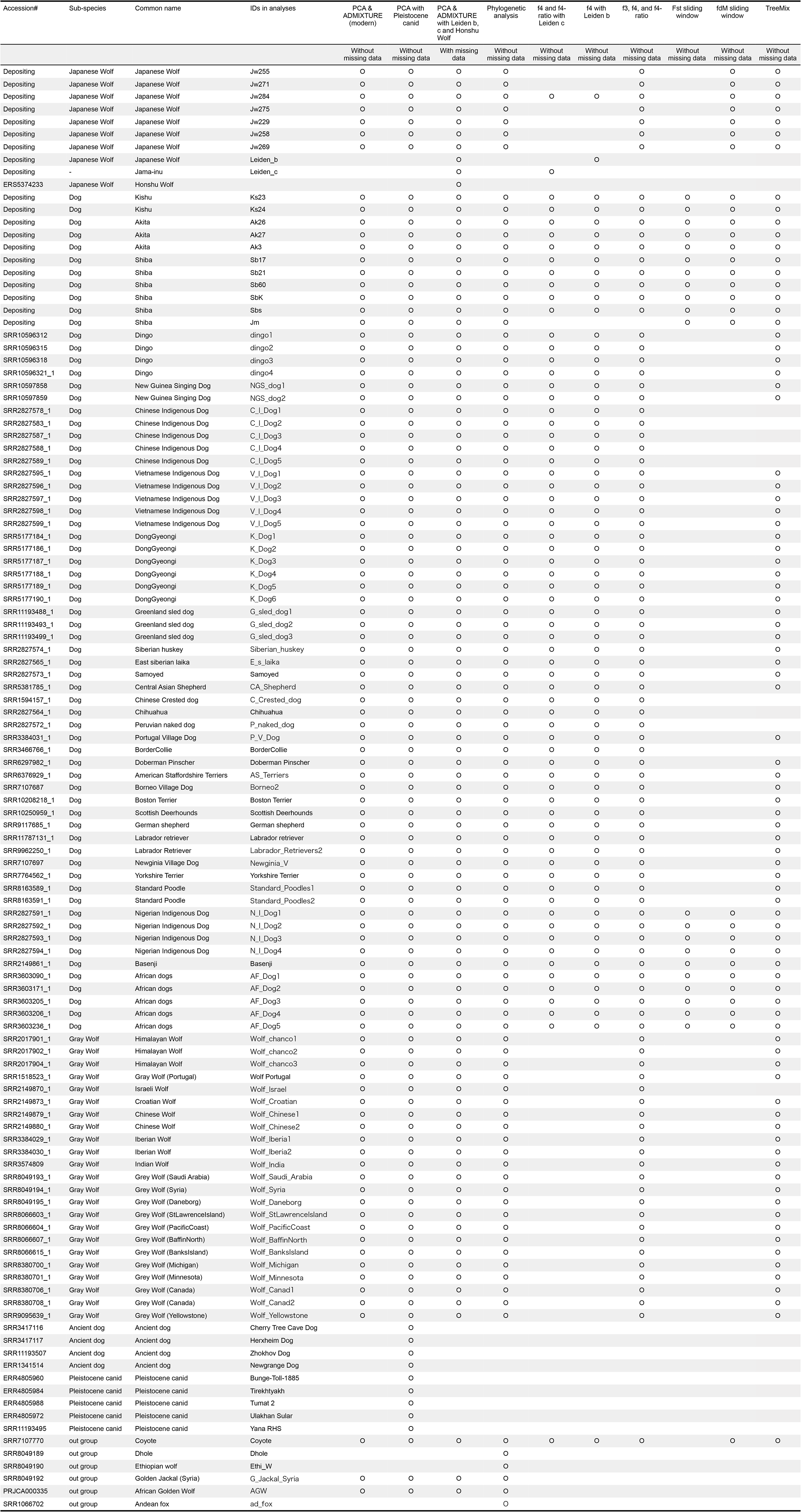
Sample information

**Table S3.** The degree of mixing of the European dog genome into the Japanese dog genome

| Japanese Dogs | *f*4-ratio (alpha) | Standard error | Z score |
| --- | --- | --- | --- |
| Akita | **0.363** | 0.019328 | 18.768 |
| Ks | **0.366** | 0.020174 | 18.133 |
| Shiba | **0.445** | 0.017633 | 25.237 |

\*European Dogs: Yorkshire\_Terrier, P\_V\_Dog, Standard\_Poodles1, Standard\_Poodles2, Scottish\_Deerhounds, BorderCollie, Boston\_Terrier, Doberman\_Pinscher, AS\_Terriers, German\_shepherd. See Table S2 for the name of the dogs.

**Table S4.** Gene flow between Japanese Wolf and ancient dogs

| Ancient Dogs | Years ago | *f4* | stderr | Zscore | *f4*-ratio(alpha) | stderr | Zscore |
| --- | --- | --- | --- | --- | --- | --- | --- |
| Newgrange | 4800 | 0.003171 | 0.000614 | **5.163** | 0.015764 | 0.00305 | **5.168** |
| Cherry Tree Cave | 5000 | 0.001028 | 0.000568 | 1.81 | 0.005231 | 0.002874 | 1.82 |
| Herxheim | 7000 | 0.003199 | 0.000606 | **5.278** | 0.016101 | 0.003041 | **5.295** |
| Zhokhov | 9500 | 0.004214 | 0.000715 | **5.897** | 0.021478 | 0.003612 | **5.947** |

**Table S5.** Genomic regions differentiated from Western Eurasian dogs and derived from Japanese Wolf

| Chromosome | 5' end | 3' end | Genes in the region |
| --- | --- | --- | --- |
| 2 | 6257975 | 6307631 | *Ankyrin repeat domain 26 (ANKRD26) |
| 3 | 36520617 | 36555078 | Upstream of MKRN3, Downstream of CHRNA7 |
| 3 | 77467500 | 77472016 | - |
| 15 | 17047771 | 17058407 | Upstream of FAM183A |
| 19 | 13299716 | 13485166 | **Upstream of INTU |
| 30 | 8023335 | 8222693 | INO80, ***DLL4, VPS18 |

\*Mice mutant in one of these genes, ANKRD26, become hyperphagia and also have enhanced adipocyte differentiation (Bera, et al. 2008; Fei, et al. 2011). The traits resulting from the mutation of this gene may have been advantageous to the diet in the Japanese archipelago during Japanese dog evolution.
\*\*INTU is part of a protein network called CPLANE, and mutations in INTU cause a number of skeletal morphological abnormalities, one of which is a flat nasal bridge (Toriyama, et al. 2016). Japanese wolves and Japanese dogs of the Jomon period (still found in a small number of Japanese dogs today) are known to have shallow stops, and perhaps mutations in this gene are related to the formation of shallow stops.
\*\*\*Notch signaling plays an important role in the development and differentiation of many cell types in diverse organisms, and DLL4, the ligand for the Notch receptor, is known to have many roles during development (Benedito and Duarte 2005). Although this gene is multifunctional and therefore traits could not be identified, it may be involved in shaping the characteristics of Japanese dogs through the Notch signaling pathway.

**Table S6.** Mapping rate (%) of reads with Japanese Wolf specific substitutions in the mitochondria DNA

| Position in CanFam3.1 | 2730 | 3346 | 4400 | 4766 | 5828 | 6461 | 7732 | 8473 | 9514 | 10464 | 10701 | 11461 | 12062 | 15225 | 15490 | Mean (%) | Estimated maximum contamination (%) |
| --- | --- | --- | --- | --- | --- | --- | --- | --- | --- | --- | --- | --- | --- | --- | --- | --- | --- |
| **Jw258** | 97.0 | 97.0 | 94.9 | 96.3 | 99.8 | 100.0 | 100.0 | 99.8 | 100.0 | 100.0 | 100.0 | 100.0 | 96.4 | 99.8 | 100.0 | 98.7 | 1.3 |
| **Jw269** | 70.3 | 80.3 | 75.0 | 76.7 | 86.7 | 100.0 | 97.9 | 83.8 | 84.2 | 89.3 | 91.0 | 84.9 | 81.8 | 86.0 | 75.0 | 84.2 | 15.8 |
| **Jw229** | 92.5 | 93.9 | 94.8 | 94.4 | 97.2 | 99.7 | 97.0 | 96.3 | 95.9 | 97.8 | 97.6 | 97.0 | 90.9 | 95.2 | 95.8 | 95.7 | 4.3 |
| **Jw255** | 95.3 | 95.9 | 95.3 | 95.4 | 97.9 | 99.9 | 98.8 | 98.3 | 97.8 | 98.1 | 98.1 | 98.5 | 96.8 | 98.0 | 97.9 | 97.5 | 2.5 |
| **Jw284** | 99.6 | 99.6 | 99.7 | 99.7 | 99.9 | 99.8 | 99.8 | 99.8 | 99.7 | 99.8 | 99.9 | 99.9 | 99.7 | 99.8 | 99.7 | 99.8 | 0.2 |
| **Jw271** | 98.5 | 98.7 | 98.5 | 98.6 | 99.2 | 99.9 | 99.4 | 99.2 | 99.1 | 99.0 | 99.2 | 99.1 | 98.9 | 99.5 | 98.3 | 99.0 | 1.0 |
| **Jw275** | 98.6 | 98.4 | 98.1 | 98.0 | 99.3 | 100.0 | 99.5 | 99.5 | 99.0 | 99.0 | 98.6 | 98.6 | 97.7 | 99.0 | 99.0 | 98.8 | 1.2 |
| **Leiden_c** | 100.0 | 98.8 | 95.6 | 100.0 | 99.2 | 100.0 | 99.3 | 100.0 | 99.2 | 99.0 | 100.0 | 100.0 | 98.9 | 100.0 | 100.0 | 99.3 | 0.7 |
| **Leiden_b** | 100.0 | 100.0 | 99.9 | 100.0 | 100.0 | 100.0 | 100.0 | 100.0 | 100.0 | 99.9 | 95.7 | 100.0 | 94.5 | 98.8 | 100.0 | 99.3 | 0.7 |

